# Recombination of ecologically and evolutionarily significant loci maintains genetic cohesion in the *Pseudomonas syringae* species complex

**DOI:** 10.1101/227413

**Authors:** Marcus M. Dillon, Shalabh Thakur, Renan N.D. Almeida, David S. Guttman

**Author notes:** These authors contributed equally to this work (Marcus M. Dillon, Shalabh Thakur). **Corresponding Author:** David S. Guttman 25 Willcocks St., ESC 4041 Toronto, ON M5S 3B2.

## Abstract

*Pseudomonas syringae* is a highly diverse bacterial species complex capable of causing a wide range of serious diseases on numerous agronomically important crop species. Here, we examine the evolutionary relationships of 391 agricultural and environmental strains from the *P. syringae* species complex using whole-genome sequencing and evolutionary genomic analyses. Our collection includes strains from 11 of the 13 previously described phylogroups isolated off of over 90 hosts. We describe the phylogenetic distribution of all orthologous gene families in the *P. syringae* pan-genome, reconstruct the phylogeny of *P. syringae* using a core genome alignment and a hierarchical clustering analysis of pan-genome content, predict ecologically and evolutionary relevant loci, and establish the forces of molecular evolution operating on each gene family. We find that the common ancestor of the species complex likely carried a Rhizobium-like type III secretion system (TTSS) and later acquired the canonical TTSS. The phylogenetic analysis also showed that the species complex is subdivided into primary and secondary phylogroups based on genetic diversity and rates of genetic exchange. The primary phylogroups, which largely consist of agricultural isolates, are no more divergent than a number of other bacterial species, while the secondary phylogroups, which largely consists of environmental isolates, have levels of diversity more in line with multiple distinct species within a genus. An analysis of rates of recombination within and between phylogroups revealed a higher rate of recombination within primary phylogroups than between primary and secondary phylogroups. We also found that “ecologically significant” virulence-associated loci and “evolutionary significant” loci under positive selection are over-represented among loci that undergo inter-phylogroup genetic exchange. These results indicate that while inter-phylogroup recombination occurs relatively rarely in the species complex, it is an important force of genetic cohesion, particularly among the strains in the primary phylogroups. This level of genetic cohesion and the shared plant-associated niche argues for considering the primary phylogroups as a true biological species.

## INTRODUCTION

*Pseudomonas syringae* is a globally significant, gram-negative bacteria that is responsible for causing a wide-spectrum of diseases on many agronomically important crops [1]. However, despite the broad host range of the *P. syringae* species complex, individual strains are highly host-specific, causing disease on only a limited range of plant species or cultivars. Furthermore, although the majority of well-characterized strains of *P. syringae* are pathogens, an increasingly number of isolates have been recovered from non-agricultural habitats that include wild plants, soil, lakes, rainwater, and clouds [2]. The diverse host range, strong host specificity, and ubiquitous distribution of *P. syringae* complex strains have made them an excellent model for studying host-pathogen interactions [3–6].

Taxonomically, the *P. syringae* species complex has been subdivided into approximately 64 pathovars based on host range and pathogenic characteristics, nine genomospecies based on DNA-DNA hybridization assays, and 13 phylogroups based on multilocus sequence analyses [7–9]. The 16S rRNA gene has also been used to differentiate strains in the *P. syringae* species complex, particularly in the context of discriminating the distinctly named species within the complex, including: *P. amygdali, P. avellanae, P. caricapapayae, P. cichorii, P. ficuserectae, P. meliae, P. savastanoi, P. syringae*, and *P. viridiflava* [10]. Nevertheless, no single locus has been found that has the ability to discriminate all *Pseudomonas* species, and importantly, these different methods often disagree on how the *P. syringae* complex should be delimited [5, 7, 8, 11–15].

Identifying genetic boundaries within and between bacterial species, and the subsequent naming of these groups, provides important insight into fundamental biological processes, as well assisting with “real world” practical decision making. From the pathologist’s perspective, who is concerned about the emergence, spread, and impact of pathogenic clones, understanding diversity and population structure is central to determining if a particular strain has the genetic potential to cause a disease on a particular crop variety and the most effective means to control the dissemination of a newly emergent pathogen clone. From a fundamental perspective, understanding natural population structure provides insight into the ecological and evolutionary pressures that give rise to natural genetic diversity, help disentangle the roles played by the different evolutionary forces, and identify specific genes that are required for the success of a strain in a particular ecological context, e.g. host specificity loci.

A significant hurdle to identifying ecologically meaningful genetic boundaries in *P. syringae* is the lack of correlation between genotypic and phenotypic similarity among strains. While *P. syringae* strains can be genetically very diverse, there are few if any definitive phenotypic traits that can reliably partition strains into major groups that are congruent with the genetic data [9, 16, 17]. For example, pathogens causing disease on a single crop are often found in multiple phylogenetic groups [8, 18–20]. Furthermore, several non-pathogenic environmental isolates are closely related to well-established *P. syringae* pathogens [21, 22]. Many of the methods that have been used to classify strains in the *P. syringae* species complex are thus forced to rely on ad hoc distinctions [23], which can lead to either the artefactual clustering of distinct lineages or splitting of cohesive monophyletic clades [24][11].

The alternative to using ad hoc distinctions or metrics to identify biological groups is to employ a theoretical framework based on evolutionary theory. Species concepts provide a theoretical basis for understanding the evolutionary and ecological forces, such as reproductive isolation, recombination, mutation, selection, and genetic drift, that drive diversification or cohesion of distinct genetic units [5]. Furthermore, unlike ad hoc species delimitation approaches, species concepts can help to define species boundaries for all isolates of a group irrespective of their specific niche or phenotype. In bacteria, the ability to horizontally exchange DNA can limit the impact of reproductive isolation; consequently, recombination, selection, and genetic drift all play a prominent role in defining species boundaries [5].

One class of models that have proven useful for understanding bacterial species are based on the concept of ecotypes. An ecotype is a genetic lineage occupying a defined niche. The basic ecotype model describes how genotypes carrying advantageous mutations arise periodically through mutation and sweep through a population as selection enables them to outcompete other members of the population [25–29]. The extent of spread of these beneficial mutations defines the boundaries of the ecotype. These recurrent selective sweeps, in combination with the accumulation of neutral mutations through genetic drift, purge genetic diversity within distinct populations, while increasing the genetic divergence between ecotypes, ultimately resulting in genetic isolation.

The primary brake on this divergence process is homologous recombination, which can transfer beneficial (as well as neutral) variation between distinct ecotypes, thus breaking down genetic isolation and maintaining genetic cohesion between ecotypes [24, 30–40]. Ultimately, the ability of recombination to disseminate these advantageous mutations among ecotypes defines the ecological boundaries of the species. The strength of recombination relative to the rate of neutral mutation and genetic drift will determine if distinct ecotypes evolve. Any decline in the frequency of homologous recombination between ecotypes, whether due to physical barriers and/or ecological partitioning, will help solidify the genetic isolation between ecotypes and formation of species. Countering this, the transfer of important genes that are critical for the exploitation of a specific niche (e.g. the interaction between a microbe and its host) may prove to be especially important for maintaining genetic cohesion in pathogenic bacterial populations like *P. syringae*.

Despite its potentially critical importance for defining species boundaries in bacteria, relatively little is known about the genome-wide extent of recombination between strains from different phylogroups of the *P. syringae* species complex because prior studies have primarily focused on a small set of housekeeping genes in the core genome [8, 41, 42]. However, we do know that at least some strains of *P. syringae* undergo relatively high rates of recombination, and this limited sample size of genes suggests that inter-phylogroup homologous recombination is considerably more rare than intra-phylogroup homologous recombination [42]. This could mean that there is no cohesive *P. syringae* species complex and each phylogroup represents a separate species. Alternatively, it is possible that the majority of inter-phylogroup recombination is occurring in the accessory genome, which would still maintain the genetic cohesion between phylogroups. It is currently not possible to distinguish between these possibilities based only on recombination analyses of a small set of core genes given that most ecologically and evolutionarily relevant genes are in the accessory genome and, by definition, only shared by a subset of strains in the *P. syringae* species complex [6, 18]. Clearly, a more thorough analysis of the rates of recombination for ecologically and evolutionarily relevant loci in the accessory genome is required to determine whether clear species barriers exist within the *P. syringae* species complex.

Here, we performed the whole-genome comparative and evolutionary analyses of 391 genomes from the *P. syringae* complex, including pathogenic isolates from diseased crops and non-pathogenic environmental isolates. In total, our collection of whole-genome sequences contains representatives from 11 of the 13 distinct phylogroups, including all seven phylogroups that we consider to be primary (1, 2, 3, 4, 5, 6, and 10) and four of the six phylogroups that we consider to be secondary (7, 9, 11, and 13). These strains enabled us to describe the phylogenetic distribution of all orthologous gene families in the pan-genome of the *P. syringae* species complex, refine the phylogenetic relationships between *P. syringae* strains using whole-genome data, predict ecologically and evolutionary relevant loci in the *P. syringae* species complex, and evaluate the impact of recombination, selection, and genetic drift on each ortholog family. Taken together, the analyses allowed us to investigate the evolutionary mechanisms that maintain genetic cohesion between *P. syringae* strains, and attain an enhanced understanding of the species barriers that exist in the *P. syringae* species complex.

## RESULTS

### Genome Assemblies and Annotations

In addition to the 135 publically available genome assemblies of *P. syringae*, we performed whole-genome sequencing and assembly on 256 new strains obtained from the International Collection of Microorganisms from Plants (ICMP) and other collaborators. The ICMP strains included 62 type and pathotype strains of *P. syringae* (BioProject Accession: PRJNA292453) [43]. Type strains are the isolates to which the scientific name of that organism is formally attached under the rules of prokaryote nomenclature. Pathotype strains have the additional requirement of displaying the pathogenic characteristics of the specific pathovar (i.e., causing specific disease symptoms on a particular host) [44]. Twenty-two non-P. *syringae* strains (twelve newly sequenced, ten from public databases) belonging to the *Pseudomonas* genus were also used as outgroups when required. In total, we analyzed whole-genome assemblies of 391 *P. syringae* strains representing 11 of the 13 phylogroups in the *P. syringae* species complex, thus enabling the most comprehensive analyses of the diversity that exists in this species to date (Supplemental Dataset S1).

All whole-genome sequencing performed in this study was accomplished using either the Illumina GAIIx platform, resulting in 36-bp or 75-bp paired-end reads, or the Illumina MiSeq platform, resulting in 152-bp paired-end reads. In sum, we generated between 614,546 to 42,765,634 paired-end reads for each genome, for an average depth of coverage ranging between 15 and 700x. Adapters and low-quality bases were trimmed from the raw reads using Trimmomatic [45], and *de novo* assembly and quality filtering were performed using CLC Genomics Workbench (CLC Genomics Work Bench 2012). After quality filtering, the final N50 value for each assembly was between 1,457 and 316,542 bps, the number of contigs was between from 59 to 5,196, and the size of each *P. syringae* genome was between 5,097,969 and 7,217,414 bps (Supplemental Dataset S2). These values represent high quality assemblies that are consistent with the draft genome assemblies that we obtained from public database (Supplemental Dataset S1; Figure S1).

*De novo* gene prediction and annotation was performed on all newly assembled and publically available genomes using a consensus approach based on Glimmer, GeneMark, FragGeneScan, and Prodigal, as implemented by DeNoGAP (Supplmental Dataset S3; see Methods) [46–50]. Reliable calls that overlapped by more than 15 bps were merged into a single coding sequence and all genes were functionally annotated by blasting against the UniProtKB/SwissProtKB database [51]. Gene ontology terms, protein domains, and metabolic pathways were also assigned to each coding sequence using InterProScan [52], while COG categories were assigned by blasting predicted genes against the Cluster of Orthologous Groups (COG) Database [53]. These methods predicted an average of 5,491 ± 25.69 (SEM) genes per *de novo P. syringae* draft assembly (Supplemental Dataset S1), and in cases where a corresponding annotation was publically available, the two annotations were largely in agreement. However, among the 135 publically available genomes, we did predict an additional 29,748 genes, for an average of 220,36 ± 11.81 (SEM) additional genes per genome (Supplmental Dataset S4).

### Evolutionary Relationships Between Strains

#### Core and accessory genetic content

Using all 413 genome assemblies (391 *P. syringae*, 22 outgroups), we clustered and differentiated homologous families using the DeNoGAP comparative genomics pipeline [46]. The 2,294,719 protein sequences present across all genomes were first clustered into 241,678 HMM families based on the stringent percent identity and alignment coverage thresholds of 70%. Similar HMM families connected via single-linkage clustering (i.e. sharing at least one sequence between the different families) were then combined, resulting in a total of 83,373 homolog families. Finally, these homolog families were split into orthologous and paralogous families using the reciprocal smallest distance approach and the MCL algorithm, resulting in a total of 98,567 ortholog families. Of the 98,567 ortholog families, 77,728 were present in at least one *P. syringae* strain, representing both the core and accessory genome content of the *P. syringae* species complex.

Despite the fact that the total number of protein-coding genes in each *P. syringae* genome is similar, the composition of each genome, with respect to the specific complement of genes, is remarkably divergent. Specifically, we estimate that only 2,457 of the 77,728 *P. syringae* ortholog families (3.16%) are part of the soft core genome, based on the presence of a given ortholog family in at least 95% of strains. This soft core genome cutoff is justified by the fact that core genome cutoffs that are overly strict result eliminate a number of genuine core ortholog families as the result of assembly and annotation errors. Indeed, as we incrementally increase the frequency of strains that a ortholog must be present in for it to be considered part of the core genome from 50% to 100%, we find that there is a sharp drop-off in the core genome size at ~95% (Figure S2), representing the point at which we expect a number of genuine core genome ortholog families to be lost due to assembly and annotation errors. The number of orthologs that are part of the hard core genome (present in 100% of strains), for example, is only 124. As more genomes are sampled, we expect the core genome size to decrease incrementally, but that this effect will diminishing as a more representative sample of the *P. syringae* complex is obtained. We asked whether we would expect further declines in the core genome size of *P. syringae* species if we sampled more genomes using a gene accumulation rarefaction curve with PanGP, which characterizes the exponential decay of the core genome as each new genome is added to the analysis [54]. The soft core genome curve plateaus as it approaches the core genome size of 2,457, when only approximately 50 genomes have been sampled (Figure 1A), suggesting that the core genome of the *P. syringae* complex would be unlikely to change significantly by sampling more *P. syringae* genomes.

**Figure 1:**
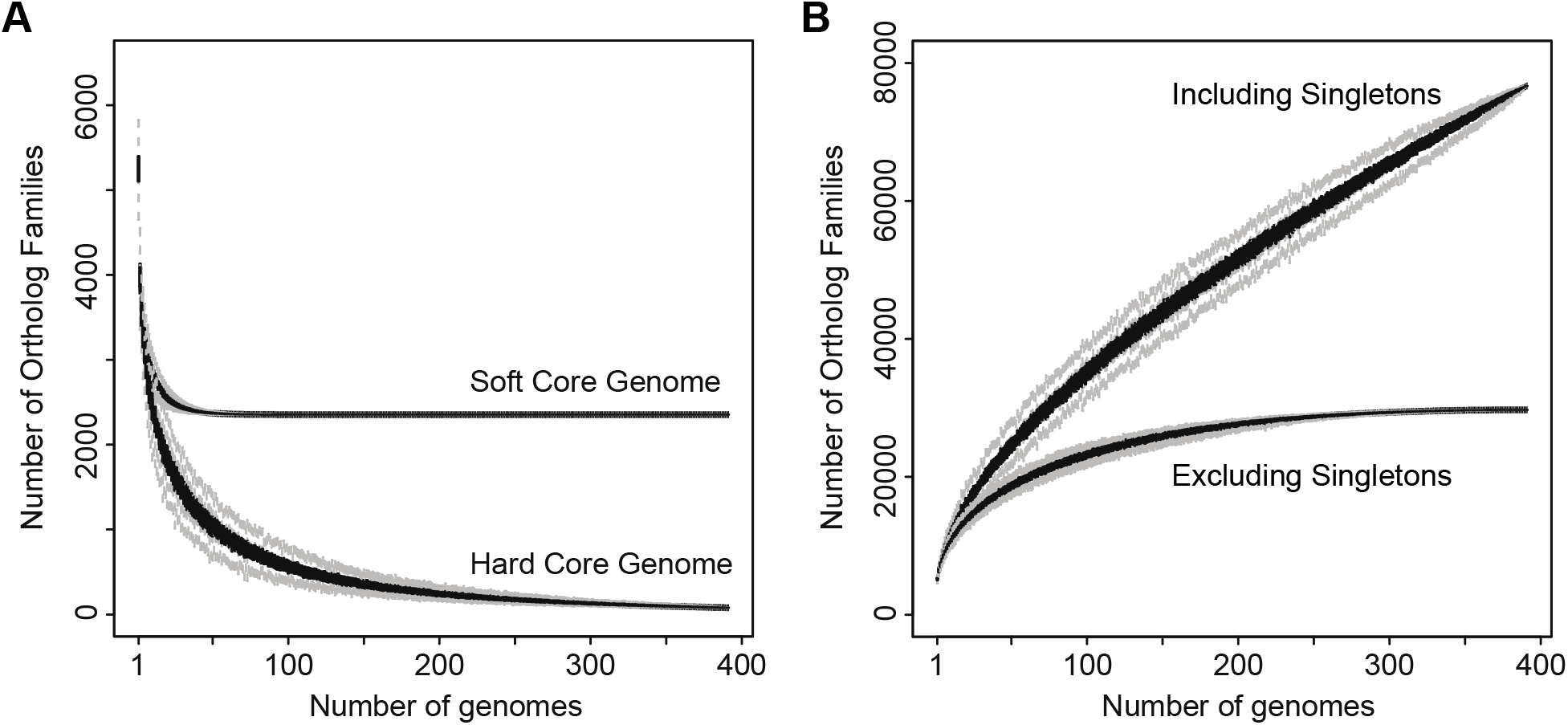
Rarefaction curves for the core (A) and accessory (B) genome of *P. syringae*, as estimated using PanGP. A) Families present in 95% (soft core genome) and 100% (hard core genome) of *P. syringae* strains exponentially decays as each new genome is added to the analysis. B) The total number of gene families identified continues to increase indefinitely as each new genome is added to the analysis when singleton gene families (families that are only present in one strain) are included, suggesting that *P. syringae* has an open pan-genome.

The small size of the core genome in the *P. syringae* species complex results in an expansive accessory genome, comprising 75,271 of the 77,728 *P. syringae* ortholog families (96.84%). Unlike the core genome, the accessory genome is expected to increase as more genomes are sampled until sufficient genomes have been sampled to capture all of the gene content diversity of the species. Only 28,165 (37.42%) of the accessory ortholog families in *P*. syringae were present in more than one strain, while the remaining 47,106 (62.58%) ortholog families were singletons present in only a single strain. We used the micropan package [55] to assess if the pan-genome of *P. syringae* is open or closed. A closed pan-genome indicates that sampling of ortholog families has neared saturation, while an open pan-genome indicates that there is still a large pool of as yet undiscovered ortholog families. Micropan estimated a decay parameter (alpha) of 0.64 using Heap’s Law Model [55], which is well below the critical threshold of alpha = 1.0 that distinguishes open from closed genomes. These findings are in agreement with a gene accumulation rarefaction analysis of the accessory genome, which has not plateaued (Figure 1B), and demonstrates that each strain introduces ~193 new ortholog families into the *P. syringae* pan-genome. Taken together, these analyses suggest that *P. syringae* possesses an open pan-genome, and that we are likely to continue to identify novel accessory ortholog families as additional *P. syringae* strains are sampled.

Overall, the distribution of ortholog families among *P. syringae* strains shows that the vast majority of families are either very common or very rare (Figure S3). This pattern is a strong indicator that lateral gene transfer is common throughout the *P. syringae* complex, and may explain its expansive accessory genome consisting of mostly singleton orthologs. While a number of these singleton orthologs were functionally annotated, signifying that they are genuine genes, 68.47% of singleton ortholog families were annotated as hypothetical proteins, compared to only 43.83% of other ortholog families (Chi-squared test; χ^2^ = 1.16 × 10^−4^, df = 1, p < 0.0001). This suggests that these genes may represent a diverse collection of yet unexplored niche specific genes in *P. syringae*, although some of these singleton ortholog families are likely the result of annotation errors associated with draft genome sequencing [56].

#### Phylogenetics

Based on multilocus sequence analysis (MLSA), the *P. syringae* species complex has currently been separated into 13 distinct phylogroups [9], seven of which we consider to be ‘primary’ phylogroups (phylogroups 1, 2, 3, 4, 5, 6, and 10) as they are monophyletic and quite genetically distinct from the more divergent ‘secondary’ phylogroups, include the traditionally recognized diversity of the species as well as nearly all of the type and pathotype strains, and predominantly carry the canonical *P. syringae* type III secretion system (discussed below). The remaining six “secondary” phylogroups (phylogroups 7, 8, 9, 11, 12, and 13) include a number of species not traditionally associated with the *P. syringae* complex such as *P. viridiflava* and *P. cichorii*, and rarely carry a canonical *P. syringae* type III secretion system. Additionally, many of the strains from the secondary phylogroups have been isolated from environmental (e.g. water and soil) sources, whereas the vast majority of strains from the primary phylogroups were isolated from aerial plants surfaces.

We first sought to refine the phylogenetic relationships between strains in the *P. syringae* species complex using a core genome alignment of the 391 strains analyzed here. The core genome tree was constructed based on a concatenated multiple alignment of the 2,457 soft core genes using FastTree with an SH-TEST branch support cutoff of 70% (Figure 2A). The core genome tree delineates these 391 strains into distinct clades representing 11 of the 13 phylogroups in the *P. syringae* species complex. Therefore, our phylogroup assignments agree with those described earlier based on a smaller collection of type strains analyzed by MLSA [7–9]. However, the clustering of strains within each phylogenetic group does differ somewhat from earlier MLSA based phylogenetic analyses [57]. This suggests that some of the more fine-scale phylogenetic relationships were not resolved, or improperly resolved due to recombination in the MLSA analysis, which were performed on a smaller collection of strains and with seven or less MLSA loci. Phylogenetic inferences based on the entire core genome should average out the majority of gene-specific biases that result from the distinct evolutionary histories of individual genes, thus providing a more accurate phylogenetic picture of the clonal relationships in the *P. syringae* species complex and enhancing our ability to explore phylogenetic relationships within and among phylogroups.

**Figure 2:**
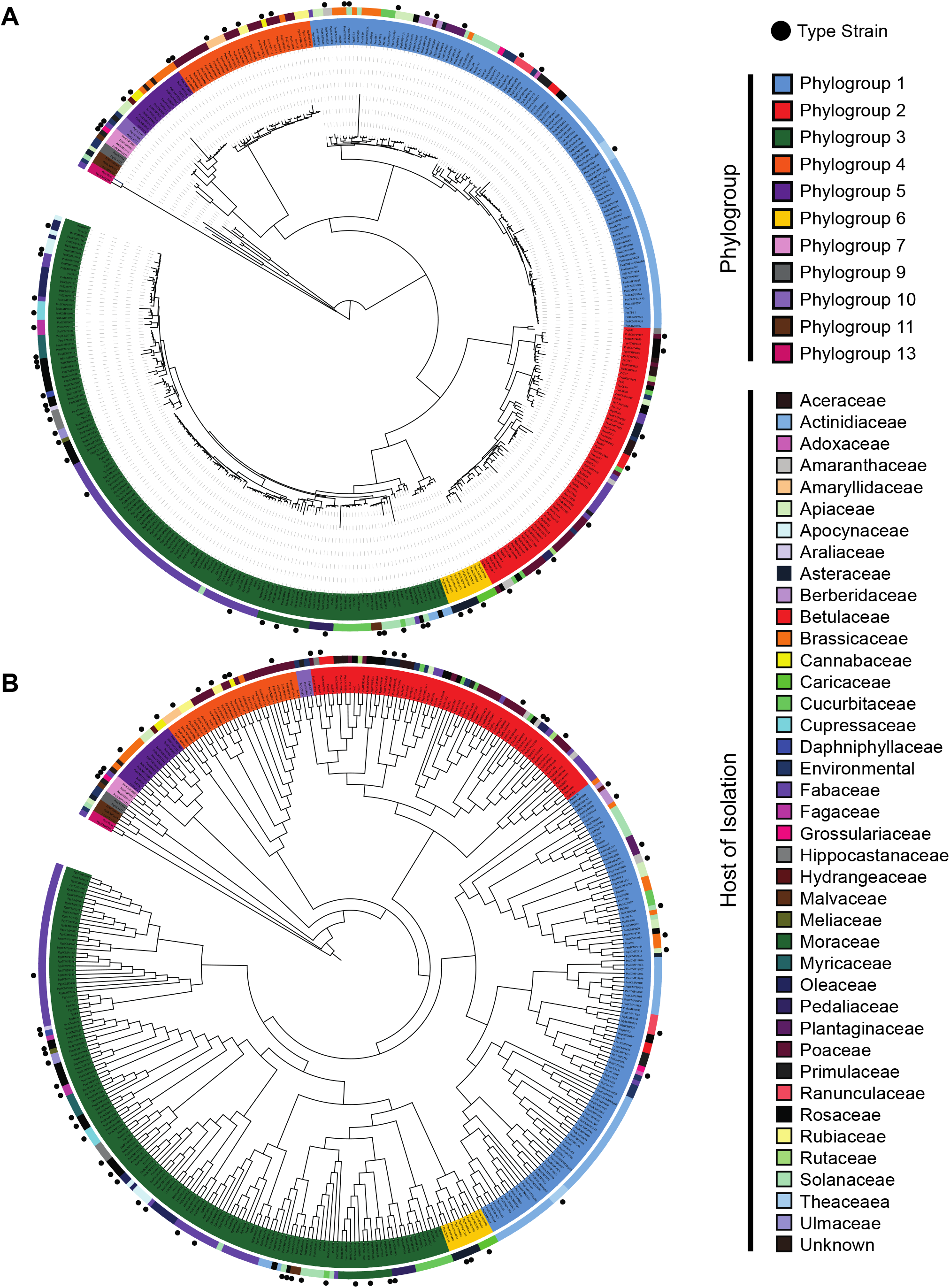
Core (A) and pan (B) genome phylogenies of *Pseudomonas syringae* strains. The core genome, maximum-likelihood tree was generated from a core genome alignment of the 2,457 core genes present in at least 95% of the *P. syringae* strains analyzed in this study. The pan-genome tree was generated by hierarchical clustering of the gene content in each strain using the Jaccard coefficient method for calculating the distance between strains and the Ward hierarchical clustering method for clustering. Strain phylogroups, hosts of isolation, and whether the strain is a type or pathotype strain are shown outside the tree.

We also assessed *P. syringae* strain relationships based on a hierarchical clustering analysis of orthologous gene content, which are simply computed as binary vectors describing the presence or absence of each ortholog family in each strain. Hierarchical clustering of the phylogenetic profiles effectively delineated *P. syringae* strains into their respective phylogroups in most cases (Figure 2B), but some key differences exist between the gene content and core genome trees. The most obvious case of incongruence between the core genome and gene content trees involves the relationship between phylogroup 2 and phylogroup 10. In the gene content tree, phylogroups 2 and 10 cluster together with all strains from these phylogroups forming a monophyletic group. This branching pattern is inconsistent with the core genome tree, where phylogroup 2 clusters with phylogroups 3 and 6, and phylogroup 10 clusters with phylogroup 5. The clustering of phylogroups 2 and 10 in the gene content tree can be traced back to their shared ortholog content. Strains from phylogroup 10 share an average of 3,918 orthologs with strains from phylogroup 2, which is more than they share with any other phylogroup, including phylogroup 5 (3,684 orthologs). There are also a number of finer scale differences between the core genome and gene content trees that involve the clustering of strains within each phylogroup. Overall, these examples of phylogenetic discordance between the core genome and gene content trees suggests that while horizontal gene transfer between strains of *P. syringae* is not sufficiently strong to consistently overwhelm the signal of vertical gene inheritance, recombination events that result in shared genome content between distantly related strains are occurring regularly between strains of the *P. syringae* species complex [58].

#### Genetic diversity

The level of divergence between phylogroups, the extremely large accessory genome, and the diversity of phenotypes within the *P. syringae* species complex has led some to propose that individual phylogroups or even specific pathovars should be considered incipient or even fully distinct species [4]. For example, Nowell et al. [58] stated that “the three *P. syringae* phylogroups [phylogroups 1, 2, and 3] are as diverged from each other as other taxa classified as separate species or even genera.” Using our expanded whole-genome dataset of *P. syringae* strains, we tested this hypothesis by quantifying the average genetic divergence between strain pairs within the same phylogroup and between strain pairs from different phylogroups. We then compared these divergence values to the pairwise divergence between three species pairs from the same genus (*Aeromonas hydrophila – Aeromonas salmonicida; Neisseria meningitides – Neisseria gonorrhoeae; Pseudomonas aeruginosa – Pseudomonas putida*), and one species pair from different genera (*Escherichia coli - Salmonella enterica*). For *P. syringae* strains, we calculated average synonymous (*Ks*) and non-synonymous (*Ka*) substitution rates across the 2,457 core genes using the “SeqinR” package in R [59]. Similarly, we calculated *Ks* and *Ka* for the distinct species pairs using 3,288 core genes for *A. hydrophila – A. salmonicida*, 1,423 core genes for *N. meningitides – N. gonorrhoeae*, 1,971 core genes for *P. aeruginosa – P. putida*, and 2,688 core genes for *E. coli – S. enterica*.

As expected, the lowest average *Ks* and *Ka* values in *P. syringae* were obtained when comparing strains within the same phylogroup, and the second lowest values were obtained when comparing strains that were from different primary phylogroups. Comparisons between *P. syringae* strains from different secondary phylogroups and of strains from primary phylogroups with those from secondary phylogroups yielded the highest *Ks* and *Ka* values, which are comparable to those that we obtained for distinct species (Figure S4). Specifically, the average *Ka* values within *P. syringae* phylogroups were all less than 0.02, and the average *Ks* values were all less than 0.20. The average *Ka* values between primary *P. syringae* phylogroups were between 0.02 and 0.04, and the average *Ks* values were between 0.30 and 0.60. With one exception, all *Ka* values between primary and secondary phylogroups, or between separate secondary phylogroups were greater than 0.05 and less than 0.10, while *Ks* values were between 0.60 and 1.00. In comparison, the *Ka* values for distinct species were 0.06, 0.15, and 0.06 for *A. hydrophila – A. salmonicida, P. aeruginosa – P. putida*, and *E. coli – S. enterica*, respectively, and their *Ks* values were 0.46, 0.74, and 0.92. The *N. meningitides* – *N. gonorrhoeae* pair was an outlier in the distinct species pairs, having a *Ka* value of 0.02 and a *Ks* value of 0.14. However, these low *Ka* and *Ks* values may be misleading because of rampant recombination between the species in this genus [60, 61]. Specifically, approximately 62.70% to 98.40% of core genes in *Neisseria* are reported to be undergoing recombination and only 1% are under positive selection [62], suggesting that the low *Ka* values in the genus are due to the elevated recombination rates that distort the molecular clock. In summary, it is clear that most *P. syringae* strains within the primary phylogroups are considerably more similar than well characterized distinct species pairs. On the other hand, most secondary phylogroups are sufficiently diverged in their core genomes to potentially warrant their separation into distinct species.

### Ecologically Significant Genes

We explored the phylogenetic distribution and diversity of what we refer to as “ecologically significant” ortholog families to better understand how these critical gene families define the ecological niche of the species complex. Specifically, we focused on any gene family previously shown to play a direct role in microbe-host or microbe-microbe interactions, such as toxins, effectors, and resistance factors. These genes included those associated with the type III secretion system (T3SS), the type III secreted effectors (T3SEs), phytotoxins, and virulence associated proteins identified using the Virulence Factors of Pathogenic Bacteria Database (VFDB) [63].

#### Type III secretion systems (T3SSs)

We investigated the phylogenetic distribution of T3SSs carried by strains in the *P. syringae* complex by searching for homologs of known proteins that constitute the structural components of different T3SSs (Figure S5). Specifically, we focused on two versions of the pathogenicity island encoding the canonical, tripartite T3SS (canonical T-PAI from *P. syringae* pv. *tomato* DC3000, alternate T-PAI from *P. viridiflava* PNA3.3a), two versions of the atypical pathogenicity island T3SS (A(A)-PAI from *P. syringae* Psy642, and A(B)-PAI from *P. syringae* PsyUB246), one version of the single pathogenicity island T3SS (S-PAI from *P. viridiflava* RMX3.1b), and one version of the Rhizobium-like pathogenicity island T3SS (R-PAI from *P. syringae* pv. *phaseolicola* 1448A) [64–72].

The canonical T-PAI T3SS is widely distributed and is found at very high frequency among strains in the primary phylogroups, and absent from the majority of strains in the secondary phylogroups (Figure 3; Figure S6). In contrast, the alternate T-PAI T3SS is only found in three strains, *Pvr*ICMP3272 and *Pvr*ICMP11296 within phylogroup 3, and *Pvr*ICMP19473 within phylogroup 7. These strains all lack the canonical T-PAI T3SS, suggesting that the alternate T-PAI acts as a replacement T3SS in these strains. Although the broad distribution of the canonical T-PAI T3SS in *P. syringae* pathogens is widely known, it is somewhat surprising that it was also present in all strains from phylogroups 9 and 10, given that these phylogroups consist of non-agricultural, environmental strains. Interestingly, some strains in phylogroup 10 have been reported to cause disease or induce a hypersensitive response (HR) in plant hosts [9], but phylogroup 9 strains have yet to be associated with any plant hosts [73]. The presence of canonical T-PAI T3SS structural genes in both of these non-agricultural phylogroups may suggest that strains in these phylogroups have the capacity to efficiently deliver effectors and cause disease in plant hosts that have yet to be examined.

**Figure 3:**
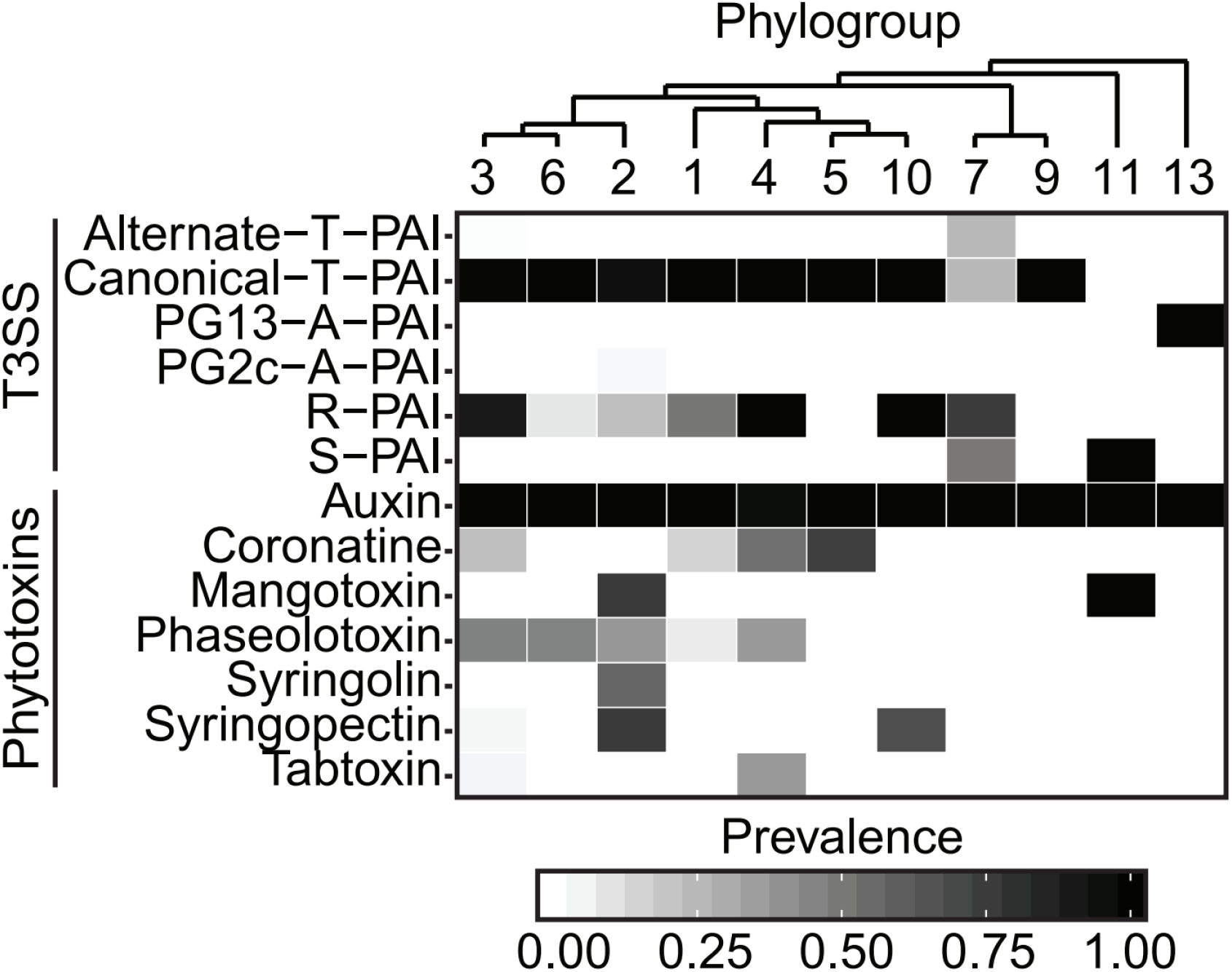
Prevalence of different forms of type III secretion systems (T3SSs) and phytotoxin biosynthesis genes in each of the *P. syringae* phylogroups. A given T3SS was considered present if all full-length, core, structural genes of the T3SS were present in the genome, while phytotoxins were considered present if more than half of the biosynthesis genes for a given phytotoxin were present in the genome.

Unlike the T-PAI T3SS, the A-PAI and S-PAI T3SSs are only present in a small subset of the *P. syringae* strains sequenced in this study. The only two homologs for the A(A)-PAI T3SS are found in phylogroup 2c, where they likely function as a replacement for the canonical T-PAI T3SS. Strains from phylogroup 2c have primarily been isolated from phyllosphere of grasses and have been widely described as non-pathogenic. However, past studies have suggested that some of these strains can efficiently deliver effectors into host cells and induce a hypersensitive response [74]. Two closely related A(B)-PAI T3SS homologs were also found in phylogroup 13. However, the A(B)-PAI T3SS in these strains is located in a different genomic region from the A(A)-PAI T3SS in strains from phylogroup 2c. Specifically, strains from phylogroup 2c contain the A-PAI T3SS between a sodium transporter and a recombination-associated protein [71], while in phylogroup 13 the A-PAI T3SS is located between a transcriptional regulator and a lytic murein transglycosylase (Figure S5). The lack of synteny between the location of the A-PAI T3SS in these two phylogroups suggests that they were independently acquired via horizontal gene transfer [69]. The S-PAI T3SS was also only identified in a small subset of the strains that we sequenced in this study, three of which are part of phylogroup 11, where they are the only T3SS in the genome, and two of which are part of phylogroup 7, where they also contain an R-PAI T3SS (Figure 3; Figure S6). Despite lacking the exchangeable and conserved effector loci (EEL and CEL, respectively) regions of the canonical T-PAI T3SS, and containing a 10kb insertion in the middle of the Hrc/Hrp cluster [67], we expect that these strains will be capable of successfully delivering effectors into some plant hosts.

The R-PAI T3SS, which closely resembles the T3SS found in *Rhizobium* species [72], is distinguished from other T3SS families based largely on the splitting of the *hrcC* gene, which codes for an outer membrane secretin protein [72]. Specifically, the *hrcC* gene is typically split into the *hrcC1* and *hrcC2* genes, separated by TPR domain (Figure S5), and in some strains, the *hrcC2* gene is split again into two additional fragments. The R-PAI T3SS is found in a large fraction of *P. syringae* strains from phylogroups 1, 2, 3, 4, 7, and 10 (Figure 3; Figure S6), but it is always present in concert with at least one other type of T3SS in *P. syringae* strains. All of these strains contain the characteristic split in the *hrcC* gene, but only seven strains, all from phylogroup 3, also contain a second split in the *hrcC2* gene. The similarity in GC-content between the *P. syringae* R-PAI T3SS genes and the rest of the *P. syringae* genome [72], the broad distribution of the R-PAI T3SS across *P. syringae* strains (Figure S6), and the ability of R-PAI HrcV protein phylogeny to effectively resolve distinct phylogroups (Figure S7) suggest that the R-PAI T3SS was likely present in the most recent common ancestor of the *P. syringae* complex. However, there is some disagreement between the inter-phylogroup relationships revealed by the HrcV protein tree and the core genome tree, with phylogroup 2 clustering with phylogroups 4 and 10 instead of phylogroup 3. This suggests that the R-PAI T3SS has also been transferred horizontally between phylogroups during the evolutionary history of the *P. syringae* species complex. From an evolutionary perspective, the presence of the R-PAI T3SS in such a large number of *P. syringae* lineages may suggest its selective benefit in nature [5], but the exact function of the R-PAI T3SS has yet to be investigated.

#### Type III secreted effector proteins (T3SEs)

The role of T3SSs is to deliver T3SEs into host plant cells to subvert the host immune response and promote bacterial growth. Therefore, we also explored the frequency and distribution of known T3SE families across *P. syringae* strains by blasting experimentally validated and predicted T3SEs against our *P. syringae* genome assemblies [75, 76]. We also attempted to identify novel T3SE candidates by searching for the universal N-terminal secretion signal and the *hrp*-box motif.

The number of known T3SE families per strain varied dramatically, from a minimum of four in strains from phylogroup 9, to a maximum of nearly 50 in some strains from phylogroup 1 (Figure 4, Figures S8). By analyzing the distribution of each effector family across *P. syringae* strains in the primary phylogroups (Figure 4), we identified three core T3SEs (*avrE1, hopAA1, hopAJ2*) that were present in some form (full-length ORF or truncated ORFs) in more than 95% of the primary phylogroup strains. Two of these core T3SEs (*avrE1* and *hopAA1*) are found in the CEL of the canonical T-PAI T3SS. In addition, a number of other T3SEs, including a third T3SE from the CEL (*hopM1*), are also broadly distributed across *P. syringae* phylogroups (Figure 4), but did not pass the core genome threshold of 95%. Interestingly, in contrast to the other T3SEs in the CEL, *hopN1* is not broadly distributed and is only found in phylogroup 1 strains.

**Figure 4:**
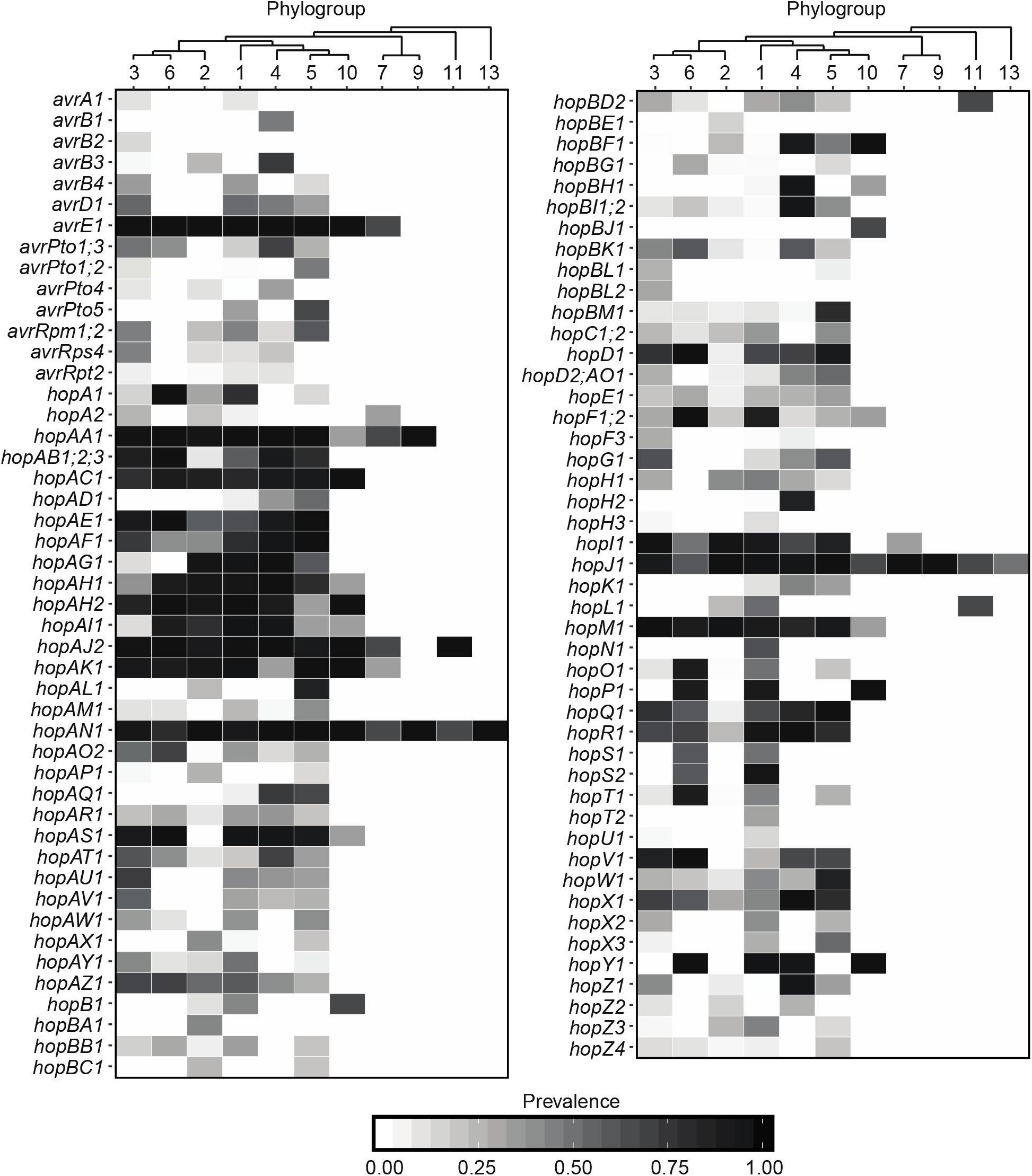
Prevalence of all known type III secreted effectors (T3SEs) in each of the *P. syringae* phylogroups analyzed in this study. T3SEs were identified using a tblastn of 1,215 experimentally verified or computationally predicted effector sequences from the BEAN 2.0 database, and were considered present if a significant hit was found in the genome (E-Value < 1^−5^). Grey scaling indicates the prevalence of each T3SE family within the respective phylogroups.

The remaining T3SEs are patchily distributed across the phylogenetic tree and a hierarchical clustering analysis of the total effector content of individual *P. syringae* strains reveals that strains from the same phylogroup can differ substantially in their T3SE content (Figure S8A). Specifically, in the T3SE content tree, phylogroup 6 strains are clustered with phylogroup 1 instead of phylogroup 3. Phylogroup 3 and phylogroup 5 strains are split in the T3SE content tree. Specifically, some phylogroup 3 strains cluster with phylogroup 1 and others cluster with phylogroup 2, while distinct clusters of phylogroup 5 strains are also found on distant regions on the T3SE content tree. Finally, while all secondary phylogroups strains, which contain considerably fewer T3SEs than primary phylogroup strains, cluster separately from primary phylogroups in the T3SE content tree, these phylogroups are often not resolved based on their T3SE contents and include the two low T3SE content strains from phylogroup 2c.

We also performed a separate analysis focusing only on variation in the exchangeable effector locus (EEL) in each of our *P. syringae* strains, which is known to be located between the *tRNA-Leu* and *hrpK1* genes. An EEL region was identified in all 380 primary phylogroup strains with the exception of the two strains in phylogroup 2c, but was only identified in four out of the eleven secondary phylogroup strains. As expected, the content of the EEL region was highly variable across strains, and a hierarchical clustering analysis of the EEL content revealed that the content of these regions does a poor job of resolving even primary phylogroup relationships (Figure S8B). For this analysis, we only included the 211 *P. syringae* strains that contained intact EEL on a single contig. Overall, the patchy distribution of T3SEs across the *P. syringae* phylogenetic tree, particularly those in the EEL, demonstrates that T3SEs are highly dynamic genes that are under frequent selection for gene gain or loss to favor adaptation to specific plant hosts and may undergo increased rates of horizontal gene transfer.

In addition to the known effector families, 6,264 additional protein sequences from the *P. syringae* species complex contained a characteristic T3SE N-terminal secretion signal and an upstream *hrp*-box promoter. We re-annotated these protein sequences using the Gene Ontology and Uniprot databases (Table S1), and found that 5,325 (85.01%) of these putative effectors were either known T3SEs that were missed in our blast similarity analysis or were sequences associated with the T3SS. The remaining 939 proteins, which were annotated with a diverse array of functions relating to metabolic processes, protein transport, signal transduction, peptidase activity, and pathogenesis, are candidates for novel T3SEs. Further computational and experimental verification of these candidate T3SEs will ultimately be required to determine if these are in fact T3SEs. However, we recommend that the 458 putative T3SEs with a *hrp*-box between 15 and 265 base-pairs from their start codons be prioritized for these studies, as has been suggested previously [77–79].

#### Phytotoxins

Phytotoxins are secondary metabolites that play a non-host-specific role in pathogenesis as well as having generalized antibacterial and antifungal properties [80]. We studied the distribution of seven well-known phytotoxin biosynthesis pathways in *P. syringae*, including auxin, mangotoxin, syringopeptin, syringolin, tabtoxin, phaseolotoxin, and coronatine by using a protein blast search of their known biosynthesis genes (Figure 3; Figure S9). Specifically, we considered phytotoxin pathways present if we identified more than half of the proteins involved in the biosynthetic pathway in a strain. Auxin appears to be the only broadly distributed phytotoxin, as genes for auxin production were found in all strains of *P. syringae* species complex, with the exception of PziICMP8959 from phylogroup 4. In contrast, mangotoxin is restricted to strains from phylogroups 2 and 11. Both syringopeptin and syringolin are also primarily restricted to strains from phylogroup 2, while tabtoxin is restricted to a small number of strains in phylogroups 3 and 4. Genes for the production of phaseolotoxin and coronatine are found in a larger proportion of phylogroups, but are still missing from many *P. syringae* strains. Overall, the majority of *P. syringae* strains only possess genes necessary to produce one or two phytotoxins; however, strains from phylogroup 2, and to a lesser extent phylogroup 4, can synthesize three or even four phytotoxins. Interestingly, phylogroup 2 strains harbor fewer T3SE genes, which suggests that phylogroups 2 strains may have evolved a unique strategy to interact with their hosts or associated microbiomes that relies more on generalized toxins as opposed to specialized T3SEs [18, 81–83].

#### Miscellaneous virulence-associated systems

Finally, we performed a search for all putative virulence factors in *P. syringae* by scanning the proteome of each strain using a BLAST search against the Virulence Factors of Pathogenic Bacteria Database (VFDB) [63]. 885 out of 17,807 orthologous protein families that were present in at least five *P. syringae* strains (4.97%) were identified as predicted virulence factors and were significantly associated with 36 different biological process (FDR p-value < 0.05) [84, 85]. These pathways included a high frequency of families involved in cellular localization, pathogenesis, flagellar movement, protein secretion, regulation of transport, siderophore biosynthesis, secondary metabolite biosynthesis, and other metabolic processes (Table S2).

### Evolutionarily Significant Genes

We explored the phylogenetic distribution and diversity of what we refer to as “evolutionarily significant” ortholog families to identify which gene families are significantly impacted by natural selection and recombination. We focused on those gene families showing genetic signatures consistent with positive selection and/or recombination. We were particularly interested in identifying loci which recombine between distinct phylogroups since these have the potential to reinforce the genetic cohesion in this diverse species complex.

#### Positive selection

We performed a codon-level analysis of natural selection using FUBAR [86] on all 17,807 ortholog families that were present in at least five *P. syringae* strains to identify families with significant evidence of positive selection at one or more residues (Bayes Empirical Bayes P-Value ≥ 0.9; dN/dS > 1). Recombination was accounted for in this analysis by using a partitioned sequence alignment and the corresponding phylogenetic tree from the output of GARD (see below), which identified 1,649 ortholog families with signatures of homologous recombination (P ≤ 0.05). A total of 3,888 ortholog families had significant evidence of positive selection at one or more codons (21.83%), with 931 of these families (23.95%) coming from the core genome and 2,957 (76.05%) coming from the accessory genome. Interestingly, this suggests that there is a significant bias for genes in the core genome to contain individual sites under positive selection (Chi-squared test; χ^2^ = 5670.60, df = 1, p < 0.0001), despite the fact that overall these genes are constrained by purifying selection and conserved across the *P. syringae* species complex.

#### Recombination

We searched for different signatures of homologous recombination in the 17,807 ortholog families that were present in at least five *P. syringae* strains using four programs: GARD [87], CONSEL [88], GENECONV [89], and PHIPACK [90]. These four methods use different underlying principles to identify recombination. GARD uses genetic algorithms to assess phylogenetic incongruence between sequence segments. CONSEL employs the Shimodaira-Hasegawa test to assess the likelihood of a dataset given one or more trees. GENECONV looks for imbalances in the distribution of polymorphism across a sequence (i.e. clusters of polymorphisms). PHIPACK calculates a pairwise homoplasy index (PHI statistic) based on the classic four gamete test [91] that assesses the minimum number of homoplasies needed to account for the linkage between two sites. Our analysis identified a total of 11,533 (64.77%) ortholog families with signatures of homologous recombination in at least one of these analyses. Specifically, GARD, CONSEL, GENECONV, and PHIPACK identified 1,616, 1,681, 4,433, and 7,379 ortholog families respectively (Bonferroni corrected P ≤ 0.05), with relatively little overlap between these packages (Figure S10). Not surprisingly, those ortholog families that displayed evidence of recombination had significantly greater average lengths (1010.09 bps ± 8.70 (SEM)) than those that did not display evidence of recombination (683.49 bps ± 10.55 (SEM)) (Welch’s Two Sample T-Test; t = 23.87, df = 14,148, p < 0.0001). This is consistent with the expectation that shorter genes are less likely to be involved in recombination because of their decreased target size and/or the decreased power of analyses of recombination on shorter genes [90, 92, 93]. The GENECONV analysis additionally classifies recombining ortholog families into intra- and inter-phylogroups recombination events, demonstrating that ortholog families that recombine within phylogroup (2,476; 55.85%) are more common than ortholog families that recombine between phylogroups (1,957; 44.15%).

Using all 11,533 ortholog families with signatures of homologous recombination, we first asked whether the well-established negative correlation between the frequency of homologous recombination and evolutionary rate could explain the reduced recombination rate between phylogroups [94, 95]. Given the wide range in strain numbers and overall diversity among phylogroups, we normalized the number of recombination events occurring between phylogroups in a number of different ways, including: recombination events per gene per strain, events per gene adjusted by branch length, events per strain adjusted by branch length, and others. The general pattern was the same regardless of the means of normalization, so we report here the analysis after normalizing recombination events per strain adjusted by branch length. Our analysis revealed a significant negative log-linear relationship between normalized recombination frequency and non-synonymous substitution rates (*Ka*) for strains within the same phylogroup and between different primary phylogroups, as predicted (Linear regression; F = 49.51, df = 30, p < 0.0001, r^2^ = 0.6227) (Figure 5A). A significant negative log-linear relationship was also observed between normalized recombination frequency and synonymous substitution rates (*Ks*) for the same strain pairs (Linear regression; F = 54.53, df = 30, p < 0.0001, r^2^ = 0.6451) (Figure 5B). In contrast, recombination events between strains from different secondary phylogroups and between strains in primary versus secondary phylogroups displayed a significant negative log-linear relationship between normalized recombination frequency and *Ka* (Linear regression; F = 10.58, df = 32, p = 0.0027, r^2^ = 0.2485) (Figure 5A). Again, this relationship was supported by comparisons of normalized recombination frequency with *Ks* for the same strain pairs (Linear regression; F = 11.40, df = 32, p = 0.0019, r^2^ = 0.2627) (Figure 5B). One of the reasons why we might not find a negative relationship between recombination rate and evolutionary rate between more distantly related strains is that other factors, like environmental isolation, are confounding recombination biases that are associated with sequence similarity.

**Figure 5:**
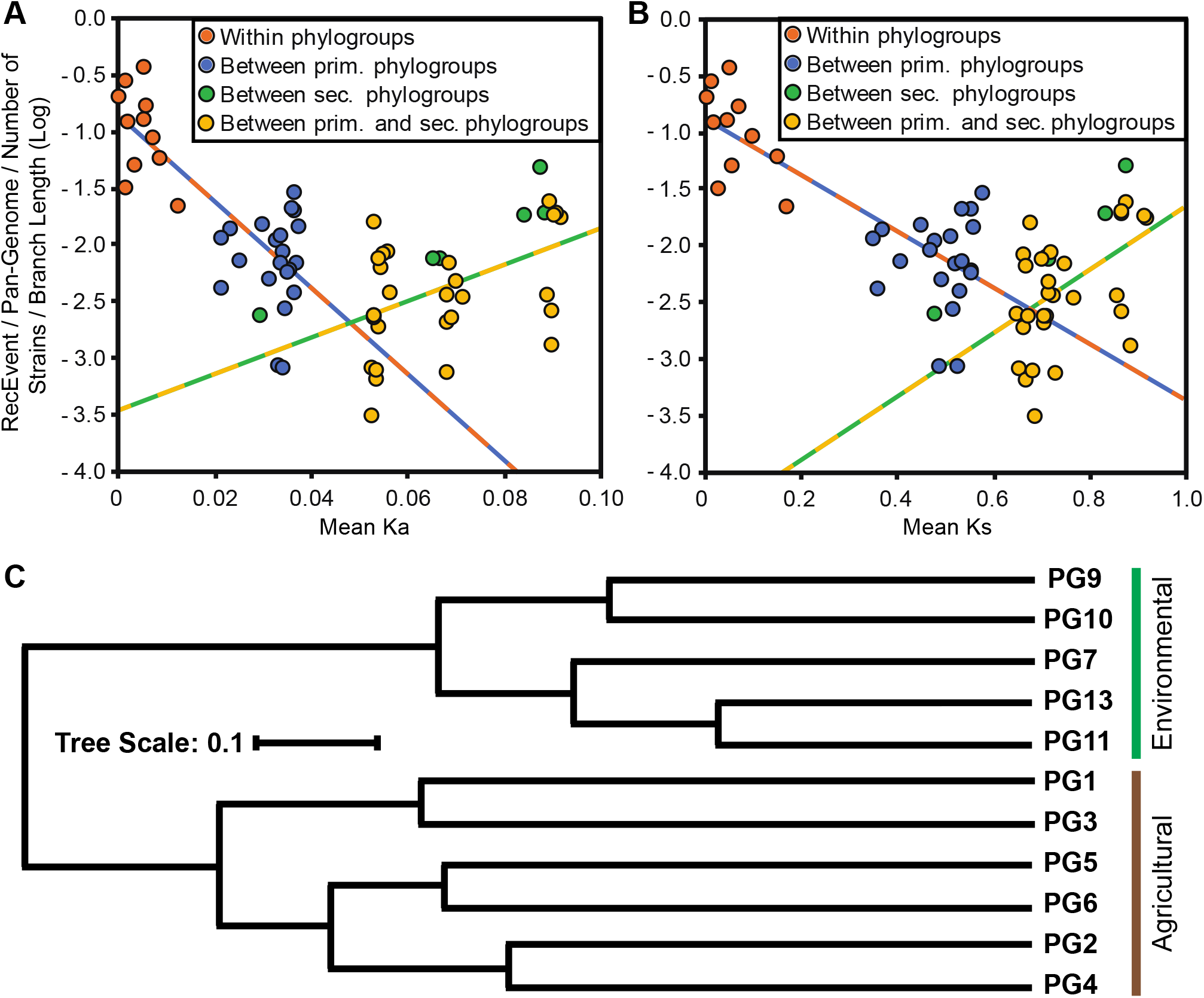
Recombination analysis between *P. syringae* strains from different phylogroups (PGs). Pairwise phylogroup recombination events were normalized based on the pan-genome size, the number of strains, and the total branch length for each phylogroups pair. A) Regression analysis of recombination rates and corresponding non-synonymous substitution rates (*Ka*). There is a significant negative log linear relationship between recombination rates and *Ka* for strains within the same phylogroup and between different primary phylogroups (F = 49.51, df = 30, p < 0.0001, r^2^ = 0.6227); however, the inverse relationship exists when comparing more distantly related strains from different secondary phylogroups and strains from primary and secondary phylogroups (F = 10.58, df = 32, p = 0.0027, r^2^ = 0.2485) B) Regression analysis of recombination rates and corresponding synonymous substitution rates (*Ks*). The same significant negative (F = 54.53, df = 30, p < 0.0001, r^2^ = 0.6451) and positive (F = 11.40, df = 32, p = 0.0019, r^2^ = 0.2627) log linear relationships were observed for strains within the same phylogroup and between different primary phylogroups, and more distantly related strains from different secondary phylogroups and strains from primary and secondary phylogroups, respectively C) Hierarchical clustering of homologous recombination frequency between phylogroups of the *P. syringae* species complex. Pairwise distances between phylogroups were calculated using the Jaccard coefficient method, based on the normalized pairwise recombination rates. Note that phylogroup 10 (PG10) is a primary phylogroup that is more closely related to phylogroups 1, 2, 3, 4, 5, and 6. Agricultural vs. Environmental labeling indicates that the bulk of the strains in these phylogroups come from these sources.

We then applied hierarchical clustering analysis to assess the relationship between phylogroups based on the frequency of recombination between them (Figure 5C) and identified two distinct clusters. One cluster contains all but one of the primary phylogroups, and therefore includes the vast majority of strains that have been isolated from agricultural environments (phylogroups 1, 2, 3, 4, 5, and 6). The second clade contains all of the secondary phylogroups, and therefore includes many strains with environmental origins (phylogroups 7, 9, 10, 11, and 13). The only exception to a clean split between primary and secondary phylogroups is phylogroup 10, which clusters with the primary phylogroups in the core genome phylogeny, but clusters with the secondary phylogroups in this analysis. This finding is interesting since two of the three strains from phylogroup 10 in our collection come from environmental sources, while the third was isolated off a non-diseased plant. These results suggest that ecological differences may also play a role in establishing recombination barriers within the *P. syringae* species complex [96]. While these relationships are robust to different methods of normalizing the number of recombination events, it is important to note that we also have much better sampling of nearly all the primary phylogroups relative to the secondary phylogroups, and therefore, much more confidence in the overall patterns of diversity found in these groups.

Previous studies have also reported significant horizontal gene transfer (HGT) between the *P. syringae* complex and other bacterial species [58]. Therefore, we performed a blastp search for all protein sequences in all 391 *P. syringae* genomes (2,176,750 sequences) against the NCBI-GenBank non-redundant protein database to identify candidate genes that have recently undergone cross-species horizontal transfer. Specifically, we considered any protein sequence with a significant match from another species in the first three blast hits to be a candidate for recent cross-species horizontal transfer. This allows us to in minimize false negatives resulting from the best matches being from the query strain or other closely related *P. syringae* strains that are present in the database. Based on these criteria, we identified 31,410 (1.44%) candidate horizontally transferred genes, and another 55,765 (2.56%) genes with no similarity matches in the non-redundant database. The most common genera involved in the putative horizontal transfer events include *Pseudomonas, Xanthomonas, Burkholderia, Klebsiella, Enterobacter, Serratia, Legionella, Pectobacterium, Pantoea, Escherichia, Salmonella, Ralstonia, Azotobacter, Achromobacter, Erwinia, Rhizobium, Bordetella*, and *Stenotrophomonas* (Figure S11A). After normalizing for the number of strains in each phylogroup, it appears as though three non-agricultural, environmentally isolated phylogroups (in rank order: phylogroups 13, 7, and 11) undergo the most HGT (Figure S11B). This finding suggests that environmental *P. syringae* strains may retain more loci obtained via HGT with other bacterial species because of increased opportunities to interact with a more diverse community of microbes, many of which could be unrelated pathogenic strains.

### Maintenance of Genetic Cohesion

In clonally reproducing bacteria, recombination is the only evolutionary process that can counter lineage diversification driven by mutation, genetic drift, and selection, thereby maintaining the overall genetic cohesion of the species. As discussed above, inter-phylogroup recombination occurs less frequently than intra-phylogroup recombination. This relationship is predicted based on the well-established log-linear relationship between sexual isolation (i.e. inverse of the recombination rate) and the level of sequence divergence due to increased difficulty of forming a DNA heterduplex as sequence divergence increases [94]. Despite this, we did find evidence that a considerable proportion of ortholog families participate in inter-phylogroup recombination, which could be an important force for maintaining genetic cohesion in the *P. syringae* species complex. We therefore wished to know the relationship between inter-phylogroup recombination and ecologically and evolutionarily significant genetic loci. Specifically, we examined whether inter-phylogroup recombination disproportionately occurred at these critical loci. To study this relationship, we classified all 17,807 orthologous gene families present in at least five *P. syringae* strains based on whether they display evidence of inter-phylogroup recombination (GENECONV), whether they were identified as ecologically significant (VFDB), and whether they were identified as evolutionarily significant (FUBAR positive selection analysis).

We first asked if there was a higher frequency of ecologically significant, virulence-associated loci among the evolutionarily significant, positively selected loci (Figure 6A). 23.50% of the 885 virulence-associated ortholog families were found to have a signal of positive selection compared to 21.75% of the 16,922 non-virulence-associated ortholog families (Chi-squared proportions test; χ^2^ = 1.58, df = 1, p = 0.2081), indicating that positive selection is not more likely to operate on virulence-associated loci in general. Second, we asked if inter-phylogroup recombination disproportionately acted on virulence-associated ortholog families (Figure 6B). 15.25% of the 885 virulence-associated families were found to recombine between phylogroups compared to only 10.77% of the 16,922 non-virulence-associated families (Chi-squared proportions test; χ^2^ = 19.08, df = 1, p < 0.0001), indicating that virulence-associated loci are significantly more likely to recombine between phylogroups than non-virulence-associated loci. Third, we asked if inter-phylogroup recombination disproportionately acted on positively selected ortholog families (Figure 6C). 13.32% of the 3,888 positively selected families were found to recombine between phylogroups compared to only 10.34% of the 13,919 non-positively selected families (Chi-squared proportions test; χ^2^ = 51.40, df = 1, p < 0.0001), indicating that positively selected loci are also significantly more likely to recombine between phylogroups than non-positively selected loci. Fourth, we asked if inter-phylogroup recombination disproportionately acted on the small set of loci that are both positively selected and virulence-associated (Supplemental Dataset S5). 20.19% of the 208 positively selected, virulence-associated ortholog families were found to recombine between phylogroups as opposed to 10.88% of the 17,599 other ortholog families (Chi-squared proportions test; χ^2^ = 17.86, df = 1, p < 0.0001). This set of orthologs include some of the most widely studied loci associated with host-microbe interactions, including numerous T3SEs, components of the flagellar system (*fliC, flg22*), phytotoxins, chemotaxis proteins, and an alginate regulatory protein (Supplemental Dataset 5). We also performed this same suite of analyses focusing exclusively on primary phylogroups (1, 2, 3, 4, 5, and 6) to examine the strength of recombination to maintain genetic cohesion in this cluster of more closely related *P. syringae* strains. Indeed, although there is still no significant correlation between ecologically and evolutionarily significant genes in the primary phylogroups, the frequency with which both ecologically and evolutionarily significant genes are transferred between primary phylogroups is even greater than it was when we considered all phylogroups (Figure S12, Table S3).

**Figure 6:**
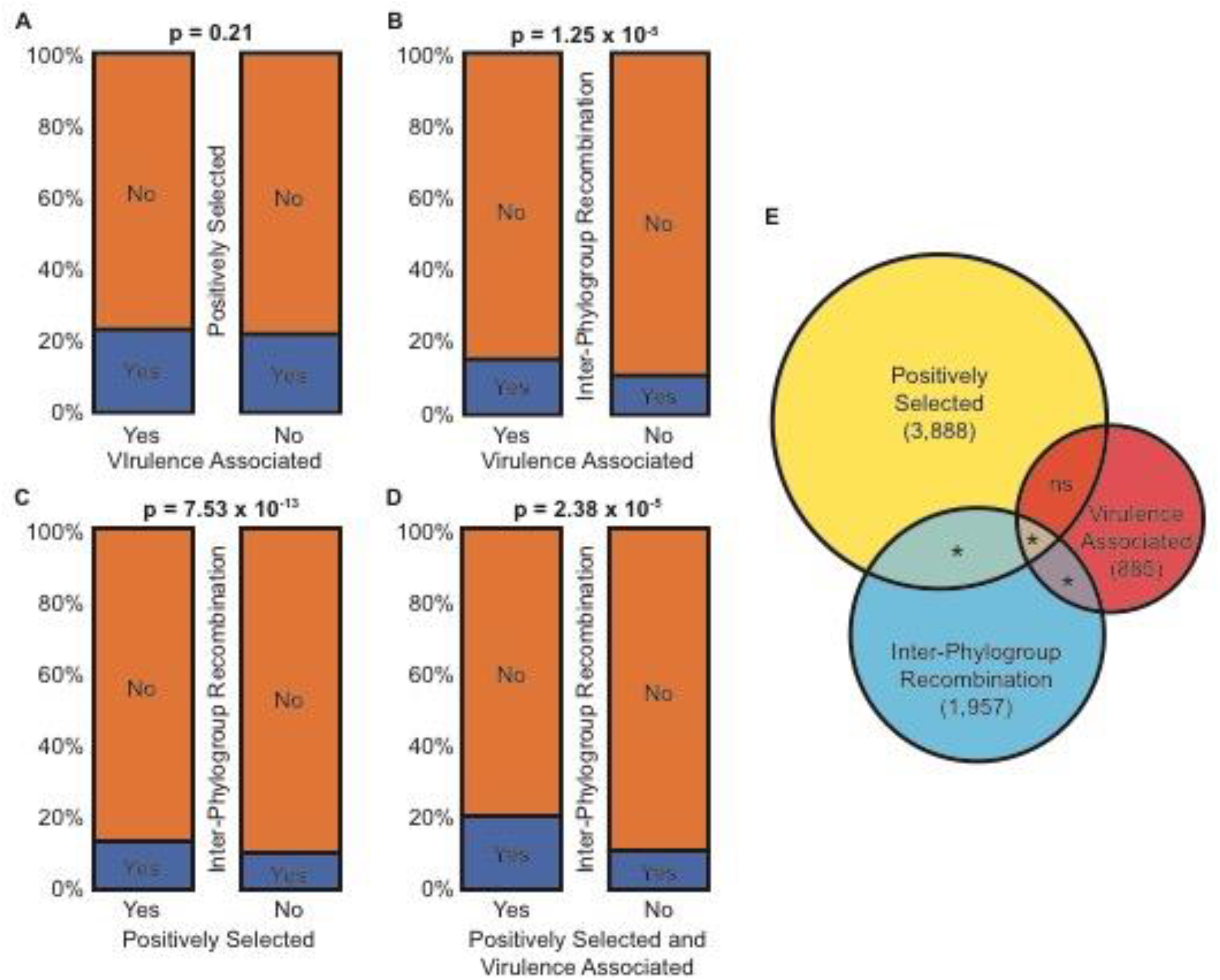
Relationships between inter-phylogroup recombination, virulence-association (“ecologically significant” loci), and positive selection (“evolutionarily significant” loci) for genes in *P. syringae* based on chi-squared proportions tests. There is no significant association between positively selected and virulence-associated genes (A). However, there is a significant positive association between gene families that have undergone inter-phylogroup recombination with virulence-associated gene families (B), positively selected gene families (C), and the small collection of gene families that are both virulence-associated and positively selected (D). The Venn diagram (E) depicts the number gene families undergoing inter-phylogroup recombination, the number of gene families that are virulence associated, and the number of gene families that are positively selected, as well as the significance of the overlap between these families.

Taken together, these results demonstrate that inter-phylogroup recombination occurs disproportionately in ecologically relevant (virulence-associated) and evolutionarily significant (positively selected) ortholog families in *P. syringae*, so while inter-phylogroup recombination may be less common than intra-phylogroup recombination, it plays a critical role in circulating genes important for maintaining the ecological niche of the species complex, and thus maintain the genetic cohesion on between all *P. syringae* strains.

## DISCUSSION

In this study, we analyzed the genomes of a diverse collection of 391 *P. syringae* strains representing 11 of the 13 *P. syringae* phylogroups to gain insight into the genome dynamics and evolutionary history of the *P. syringae* species complex. We reveal that *P. syringae* has a large and diverse pan-genome that will likely continue to expand with the sampling of more strains. We also demonstrate strong concordance at the phylogroup level between the refined core genome and gene content trees of *P. syringae* strains with a few exceptions, suggesting that while horizontal gene transfer between *P. syringae* phylogroups is typically insufficient to distort the phylogenetic signal from vertical inheritance of gene content, there are cases where it has distorted relationship among subgroups. Furthermore, by investigating the distribution of ecologically and evolutionary relevant loci in the *P. syringae* species complex and the rates of intra-and inter-phylogroup recombination of these genes, we also demonstrate that despite its relative rarity, inter-phylogroup recombination is a critical cohesive force that disproportionately facilitates the spread of ecologically and evolutionarily significant loci across *P. syringae* phylogroups.

### Core and Accessory Genetic Content in the *P. syringae* Pan-Genome

The *P. syringae* pan-genome is vast and extremely diverse, comprising a total of 77,728 ortholog families. Yet, very few of these ortholog families are present at high frequency in the *P. syringae* species complex. A rarefaction analysis demonstrates that the composition and size of core genome stabilizes after sampling approximately 50 strains at ~2500 genes. This is slightly smaller than estimates from three prior studies that identified core genome sizes of 3,397 [18], 3,364 [58], and 3,157 [97]. However, these prior studies were mostly restricted to the primary phylogroups, and only the Mott *et al*. study [97] was performed with more than 50 strains. The *P. syringae* core genome size is also comparable to the core genome sizes of other pathogenic Proteobacteria, including: *P. aeruginosa* (2,503) [98], *Erwinia amylovora* (3,414) [99], and *Ralstonia solanacearum* (2,543) [100]. This raises the possibility that different pathogenic bacteria may have similar core metabolic requirements; however, the extent to which the core genome content is conserved across species will require further investigation.

Our analysis further clarifies and expands our understanding of the highly dynamic nature of the *P. syringae* accessory genome. The gene family size distributions (Figure S3) suggest that a relatively small number of gene families are found in more than ten strains (16.36%), while the majority of families (62.58%) are only found in a single strain. The pan-genome rarefaction curve (Figure 1B) demonstrates that the pan-genome of *P. syringae* remains open after sampling 391 strains, and will therefore continue to increase in size as more diverse *P. syringae* strains are added to the analysis at a rate of ~193 new ortholog families for each new strain analyzed. The tendency of gene families to be present in only a single strain is often attributed to a species’ ability to acquire novel DNA through horizontal gene transfer [101]. However, the ubiquitous distribution of *P. syringae* strains across the globe is likely also a key contributor to the diverse gene content of different strains, as many strain-specific genes may be under selection only in specific environments. A large number of the strain-specific gene families that were identified in this study are annotated as hypotheticals with no similar sequences in the sequence databases, and thus may represent a diverse collection of niche specific genes in *P. syringae* that are entirely unexplored. However, as we have already acknowledged, it is also important to recognize that some of these strain specific genes may be artefactual due to sequencing and assembly errors [56]. Furthermore, although the *P. syringae* pan-genome remains open, we believe we have sampled the majority of higher-frequency genes since our rarefaction analysis on non-singleton orthologs did plateau (Figure 1B).

### Phylogenetic Relationships and Diversity Among *P. syringae* Strains

Investigating the relationship between core genome and gene content trees can shed important insight into the lifestyle and evolutionary history of bacterial species. Specifically, strong discordance between core genome and pan-genome trees is suggestive of extensive genomic flux among lineages [102], which obscures the clonal relationship between strains in the gene content tree. For example, genome analyses of core genome and gene content in the marine bacteria *Vibrio* have shown strong discordance, suggesting extensive horizontal transfer between lineages [103]. However, other species like the marine bacterium *Prochlorococcus* have concordant core genome and gene content phylogenies [104], suggesting that horizontal transfer has played a lesser role in their evolutionary history.

In *P. syringae*, the core genome and gene content trees are largely concordant at the level of phylogroups. The one major exception to this concordance is the relationship between phylogroups 2 and 10, which cluster more closely in the gene content tree than they do in the core genome tree. Previous studies have shown that phylogroups 2 and 10 have similar virulence repertoires [57], and that almost all strains from these phylogroups have high ice nucleation activity [9, 73, 105]. This elevated gene content and phenotypic similarity likely reflects similarity in the lifestyles and ecology of strains from these phylogroups, which may be the result of increased horizontal transfer, convergent evolution, or both. Indeed, we find that the 2,832 gene families that are in the soft-core genome (>95% of strains) of both phylogroups 2 and 10 are significantly more likely to be evolutionarily significant (Chi-squared proportions test; χ^2^ = 832.31, df = 1, p < 0.0001) and ecologically significant (Chi-squared proportions test; χ^2^ = 9.72, df = 1, p = 0.0018) than the remaining 14,975 non-core families. However, gene families in the soft-core genome of phylogroups 2 and 10 are significantly less likely to be involved in inter-phylogroup recombination events than other genes (Chi-squared proportions test; χ^2^ = 15.22, df = 1, p < 0.0001). This suggests that phylogroups 2 and 10 strains do not exchange more genes than the rest of the *P. syringae* species complex through recombination. Consequently, convergent evolution likely plays a key role in the increase of shared genes between these two phylogroups. It is nevertheless important to emphasize that the *P. syringae* core genome and gene content trees are largely concordant at the level of phylogroups, which suggests that although we do find some evidence of genomic flux, the rate of inter-phylogroup horizontal transfer is not sufficient to obscure the phylogenetic signature of vertical gene inheritance.

The *P. syringae* species complex is unquestionably highly diverse, and claims have been made that the diversity between phylogroups is actually greater than the observed diversity between well-established species [58]. We used the entire soft core genome alignment to estimate the level of genetic divergence between all phylogroups to explore whether distinct phylogroups do in fact have consistently higher genetic divergence than distinct species pairs (Figure S4). We determined that average *Ka* and *Ks* values among strains in the primary phylogroups were less than the average values between *P. aeruginosa* and *P. putida* strains, and *E. coli* and S. *enterica* strains. The average among primary phylogroup *Ka* value was also lower than the average values between strains of *A. hydrophila* and *A. salmonicida*, although the *Ks* value was roughly similar. Estimates of *Ka* and *Ks* between *N. gonorrhoeae* and *N. polysaccharea* are considerably lower than those of both *P. syringae* phylogroups and other distinct species pairs, but the *Neisseria* genus is known to be highly recombinogenic, which can distort evolutionary rates, making this species pair a likely outlier [62]. In contrast, both the average *Ka* and *Ks* values obtained when comparing strains between primary and secondary phylogroups, and those between secondary phylogroups are more consistent with the distinct species pairs, with a few exceptions. Overall, these analyses suggest that the primary phylogroups are not excessively divergent relatively to other bacterial species, in contrast to the secondary phylogroups, which may be sufficiently divergent to be considered distinct species.

### Phylogenetic Distribution Ecologically Significant Genes

A unifying feature among all strains in the *P. syringae* species complex is the presence of at least one T3SS. The most common T3SS in the *P. syringae* species complex is the canonical T-PAI T3SS, and consistent with prior studies, we found that nearly all agriculturally associated strains carry one. In addition, we also found that a number of non-agricultural strains from phylogroups 9 and 10 possess a canonical T-PAI T3SS. These data are consistent with an earlier report of the presence of a canonical T-PAI T3SS in non-agricultural strains from phylogroup 1A [21, 22], some of which were shown to cause disease on tomato. Although the host-range of these non-agricultural strains from phylogroups 9 and 10 has yet to be studied experimentally, it raises the interesting possibility that they may be pathogens of wild plant species and act as a reservoir for the recurrent emergence of crop pathogens.

In addition to the canonical T-PAI T3SS, we also found that many *P. syringae* strains possess an R-PAI T3SS, while the A-PAI and S-PAI T3SSs are found in a small number of strains isolated in discrete phylogroups. The A-PAI and S-PAI T3SSs are always present in the absence of the canonical T-PAI, suggesting that they may serve as a replacement T3SS in a different niche. In contrast, the R-PAI T3SS is always present in concert with at least one other T3SS. Bacteria with multiple T3SSs that have complementary functions have been reported previously [106, 107]. For example, *Salmonella* species contains two different T3SSs known as SPI-1 and SPI-2 [106]. SPI-1 promotes bacterial pathogenicity by facilitating host invasion, while SPI-2 is critical for survival, replication and dissemination of the bacteria after it enters the host cell [108]. This is also not the first study report of the presence of the R-PAI T3SS outside of *Rhizobium* species. A wide array of symbiotic and non-pathogenic bacteria, including *Photorabdus luminescens, Sodalis glossindicus, Pseudomonas fluorescens*, and *Desulfovibrio vulgaris*, have also been reported to harbor the R-PAI T3SS [108]. Although its expression in *P. syringae* is low and its function outside of *Rhizobia* remains unclear [72], the broad distribution of this the R-PAI T3SS across *P. syringae* strains implies that it is likely of functional importance for a number of strains in the complex.

The phylogenetic distribution of the different T3SSs and our phylogenetic analysis of the conserved HrcV protein from all T3SSs also sheds critical light on the evolutionary history of each T3SS in the *P. syringae* species complex. The broad phylogenetic distribution of the T-PAI T3SS has led some previous studies to conclude that it was present in the most recent common ancestor of the *P. syringae* species complex [109, 110], while others have suggested that the canonical T-PAI may have been acquired after the divergence of the primary and secondary phylogroups [69, 73]. Indeed, the patchy distribution among strains in the secondary phylogroups (i.e. found in only 37.50% of secondary phylogroup strains vs. 97.91% for primary phylogroup strains) observed here provides evidence that the canonical T-PAI was acquired after the divergence of the primary and secondary phylogroups. However, acquisition by the common ancestor of all *P. syringae* and subsequent loss by secondary phylogroup lineages is also a possibility.

Two additional lines of evidence support the early acquisition of both the T-PAI and the R-PAI T3SSs. First, the genomic region encoding these T3SSs shares the same %GC as the rest of the genome [6, 72]. Second, the HrcV genealogies from both the T-PAI and the R-PAI T3SSs are generally congruent with the core genome tree (Figure 2A; Figure S6), indicating a common evolutionary history. In contrast, the rarity of the A-PAI and S-PAI T3SSs in the *P. syringae* complex suggest later horizontal transfer into only a few *P. syringae* lineages. Specifically, the A-PAI T3SS appears to have been acquired independently in phylogroup 13 and a small group of phylogroup 2 strains (phylogroup 2c), as evidenced by the unique location of the A-PAI T3SS in these two genomes. The S-PAI T3SS, which is most closely related to the T3SS found in *Erwinia* and *Pantoea* species, is also present in two distantly related phylogroups (7 and 11) which are reported to be pathogenic on some plants [9].

As shown in previous studies [6, 18, 111], T3SEs that are delivered by the T3SS are patchily distributed across the *P. syringae* species complex with a few exceptions. The presence of these T3SEs in only a small but diverse suite of strains suggests that horizontal gene transfer is common in these families and that they are subject to strong diversifying selection. Specifically, T3SEs are known to experience frequent gain/loss events and rapid sequence diversification to obtain new functional capabilities or avoid host immune recognition [18, 112–114]. The phylogenetic distribution and diversification of the effectors analyzed in this study suggests that both of these evolutionary forces are at play in a large number of the *P. syringae* T3SE families. Despite the patchy distribution of most T3SEs, prior studies have identified a set of four core T3SEs, which include *avrE1, hopAA1, hopM1*, and *hopI1* (Lindeberg *et al*., 2012; O’Brien *et al*., 2011a). We confirmed this characterization for the *avrE1* and *hopAA1* families, but the *hopM1* and *hopI1* effectors are not present in more than 95% of the strains analyzed in this study, even though they are present in the majority of strains from the primary phylogroups. In addition to *avrE1* and *hopAA1*, we also identified a third core T3SE, *hopAJ1*, and two other T3SE families, *hopAN1* and *hopJ1*, that are present at some frequency in all eleven phylogroups. Finally, using an HMM-modelling approach that searches for the conserved N-terminal secretion signal and the hrp-box promoter of known T3SEs, we have also proposed a new set of novel T3SEs in the *P. syringae* species complex that are strong candidates for functional assays (Table S1).

### Recombination and Genetic Cohesion in the *P. syringae* species complex

Recombination plays a significant role in the evolution of bacteria [95, 115], and while it can lead to either genetic diversification or homogenization depending on the population structure of the donor and recipient strains, the latter role is particularly important in maintaining genetic cohesion within a species [31, 34, 115, 116]. Previous studies in *P. syringae* have reported that recombination between phylogroups is relatively rare [8, 58, 117]. However, these studies were based on analyses of a small set housekeeping genes in a limited collections of strains, so lacked a sufficient genomic and sampling depth to draw firm conclusions about the extent of recombination across the pan-genome. This is particularly important because it has been suggested that horizontal transfer occurs at a relatively high rate in the accessory genome and has a disproportionate effect on strain adaptation in nature [5, 18, 58]. Our analysis found a signature of recombination in 11,533 (64.77%) of the 17,807 ortholog families that were present in at least five *P. syringae* strains. Among the 4,433 recombination events identified by GENECONV, 2,476 (55.85%) of these events were intra-phylogroup recombination events, while the remaining 1,957 (44.15%) were inter-phylogroup recombination events. These findings reaffirm that recombination within phylogroups is more common than recombination between phylogroups, likely as a result of the well-established linear relationship between sequence divergence and the logarithm of the recombination rate [94, 95]. However, while sequence similarity appears to be the key factor determining the rate of recombination between relatively closely related strains within the primary phylogroups, our data suggest that recombination between more distantly related strains appears to be governed by other forces (Figure 5). A particularly intriguing finding is that phylogroup 10 strains cluster with secondary phylogroup strains with respect to their pairwise recombination frequency, despite the fact that phylogroup 10 is a primary phylogroup in the core genome tree (Figure 2). The major distinction between phylogroup 10 strains and the bulk of the primary phylogroup strains is that they were isolated from non-agricultural sources, as were most of the secondary phylogroup strains. This may indicate that ecology plays a more important role in determining the extent of recombination than sequence similarity, at least for long-distance (e.g. between phylogroup) genetic exchange.

Although inter-phylogroup recombination is rarer than intra-phylogroup recombination overall, we also used our expanded dataset to explore whether specific evolutionarily and ecologically important gene families more frequently undergo inter-phylogroup recombination than other gene families. For ecologically important genes, we used all virulence associated orthologous gene families that were identified by the VFDB (885/17,807; 4.97%). For evolutionarily important genes, we used all orthologous gene families determined to be positively selected at least one site by FUBAR (3,888/17,807; 21.83%). The analysis showed that both ecologically and evolutionarily important gene families are more likely to be subjected to inter-phylogroup recombination than other gene families (Figure 6). This finding is consistent with the observation that ecologically adaptive genes are successfully transferred at high rates among diverse strains in a species complex [118], and suggests that inter-phylogroup recombination disproportionally spreads ecologically and evolutionarily important genes across phylogroups, which may help maintain genetic cohesion within the *P. syringae* species complex.

### Fundamental Evolutionary Principles for Delimiting *P. syringae* Species

There is a long history to the debate over the appropriate way to delimitate species within the *P. syringae* complex [12], stemming from the use of largely arbitrary and ad hoc species delimitation cutoffs in DNA-DNA hybridization assays, MLST analyses, and pathotype designations [12, 17, 119, 120]. Importantly, these prior studies have largely been poorly-powered in terms of both the number of strains and the number of genes analyzed. Because the current study dramatically increases both the number and diversity of *P. syringae* strains sampled, we obtain a unique perspective into the ecological and evolutionary forces operating in the *P. syringae* species complex, and suggest that future work to delimit the complex should be founded in fundamental evolutionary processes.

From an ecological perspective, species differentiation results from the adaptation of two or more subpopulations to different environments or niches [96, 121]. Here, diversifying selection among a few loci that are essential for differential adaptation to alternative environments can drive speciation in the absence of barriers to recombination. There is evidence that this has occurred in *P. syringae*, given the broad global distribution and diverse disease-causing capabilities of *P. syringae* strains [1]. Specifically, Moteil *et al*. show weak ecological differentiation between an agricultural pathogenic *P. syringae* population and a closely related environmental population of *P. syringae*, despite there being no barrier to recombination between these populations [22]. However, it is currently unclear what the differentially selected loci in these populations are and whether they have sufficiently diverged to be considered an early speciation event. Furthermore, the lack of correlation between the core genome phylogenetic profile of *P. syringae* strains and their pathovar designations suggests that there are many different pathways for adaptation to a single host, so ecological differentiation on its own is likely a poor way to speciate the *P. syringae* species complex [9, 18, 21, 22]. Future studies should focus on expanding the dataset of non-agricultural *P. syringae* strains so that we can more effectively distinguish and analyze loci that are differentially selected in ecologically divergent strains.

Both sequence clustering and recombination barriers have been used to delimit bacterial species based on evolutionary principles [122]. Yet, even with the growing abundance of genomic data, it is unlikely that any one criteria will adequately resolve species barriers in the *P. syringae* complex, largely due to the fluid nature of bacterial genomes. However, given what we now know about the phylogenetic relationships between strains, the distribution of ecologically and evolutionarily important genes, the disproportionately high rate of inter-phylogroup recombination among ecologically and evolutionarily significant loci, and finally, the common ecology of diverse *P. syringae* strains, we propose that there is no ecologically or evolutionarily justifiable basis to split the strains of the primary phylogroups of *P. syringae* into separate species. In fact, *P. syringae* provides an outstanding example of how recombination, despite being relatively infrequent, maintains genetic cohesion is this very widespread, diverse, and globally significant lineage.

## METHODS

### Genome Sequencing and Assembly

A total of 391 *P. syringae* strains and 22 outgroup *Pseudomonas* strains were used in this study (Supplemental Dataset S1). The genome assemblies and annotations for 145 of these strains were obtained from public sequence databases, including NCBI/GenBank, JGI/IMG-ER, and PATRIC [123–125]. The remaining 268 strains were obtained from the International Collection of Microorganisms from Plants (ICMP) and other collaborators, and were sequenced, assembled, and annotated in the Center for the Analysis of Genome Evolution and Function (CAGEF) at the University of Toronto. For these strains, DNA was isolated using the Gentra Puregene Yeast and Bacteria Kit (Qiagen, MD, USA). Purified DNA was then suspended in TE buffer and quantified with the Qubit dsDNA BR Assay kit (ThermoFisher Scientific, NY, USA). Paired-end libraries were generated using the Illumina Nextera XT DNA Library Prep Kit following the manufacturer’s instructions (Illumina, CA, USA), with 96-way multiplexed indices and an average insert size of ≈400 bps. All sequencing was performed on either the Illumina MISeq or GAIIx platform using V2 chemistry (300 cycles). Following sequencing, read quality was assessed with FastQC [126] and low-quality bases and adapters were trimmed using Trimmomatic v0.30 (ILLUMINACLIP: TruSeq3-PE.fa, Seed Mismatch = 2, Palindromic Clip Threshold = 30, Simple Clip Threshold = 10; SLIDINGWINDOW: Window Size = 4, Required Quality = 15; LEADINGBASEQUALITY = 3; TRAILINGBASEQUALITY = 3; MINLEN = 25) [45].

The trimmed paired-end reads for each of the 268 *Pseudomonas* genomes sequenced at CAGEF were *de novo* assembled into contigs using the CLC assembly cell v4.2 program from CLCBio (Mode = fb, Distance Mode = ss, Minimum Read Distance = 180, Maximum Read Distance = 250, Minimum Contig Length = 200). All contigs that were less than 200 bps long were then removed from each assembly and the raw reads from each strain were re-mapped to the remaining contigs using clc_mapper. Next, using clc_mapping_info and clc_info, we calculated the read coverage for each contig in each assembly and compared that with the average contig coverage of the genome assembly to identify contigs with atypical coverage (> 2 standard deviations from the average contig coverage). These atypically covered contigs were then compared to the EMBL plasmid sequence database and the GenBank nucleotide database using BLAST and were removed from the assembly if they were not identified as part of a plasmid sequence.

Gene prediction for these 268 draft *Pseudomonas* assemblies was performed using DeNoGAP [46], which predicts genes based on the combined output of Glimmer, GeneMark, Prodigal, and FragGeneScan [47–50, 127]. For most genes, these algorithms accurately predicted both the start and the stop positions, but in some instances, genes were incomplete (missing appropriate start and/or start coding). In these cases, we extended the gene as a triplet codon until a stop codon was found at both the 5’ and 3’ end. The first Methionine codon downstream from the 5’ stop codon was considered the start codon, while the first 3’ stop codon was considered the stop codon. This approach allowed us to obtain complete coding sequences for a number of incomplete genes, but for others we were unable to predict a start and stop codons due to a contig break or an assembly gap. These and any other genes that contained runs of N’s were considered partial genes and were excluded from the final dataset to avoid complications in downstream comparative and evolutionary analyses. Furthermore, complete coding regions that were only predicted by one program and could not be verified by blasting against the UniProtKB/SwissProt database or pass a minimum length cutoff of 100bps were discarded. The final collection of coding sequences was then sorted by genome location, and any coding regions that overlapped by more than 15 bases were merged into a single coding sequence.

All complete genes were then annotated using a blastp search of the corresponding protein sequences for each gene against the UniProtKB/SwissProt database with an e-value threshold of 1^−5^ [51]. The name and/or description of the best hit was assigned to the corresponding protein and proteins that did not have any significant hits were assigned as hypothetical proteins. Gene ontology terms, protein domains, and metabolic pathways were also annotated in each complete gene using InterProScan v.5 (E-Value < 1^−5^) [52]. All complete genes were also assigned Cluster of Orthologous Group (COG) categories using a blastp search against the COG database (E-Value < 1^−5^) [53, 128]. However, COG families were only assigned if the protein query had high sequence identity and coverage (> 70%) with at least three member sequences in the family.

### Ortholog Prediction and Phylogenetic Analysis

We clustered all complete protein sequences from the 413 *Pseudomonas* genomes described above, which included 391 *P. syringae* strains representing 11 of the 13 phylogroups, into putative homolog and ortholog families using DeNoGAP [46]. First, all protein sequences from the closed genome of *P. syringae* DC3000 were used to construct seed HMM families for DeNoGAP [65], using an all-vs-all pairwise protein sequence comparison with phmmer (E-Value < 1^−10^) [129]. Proteins that had greater than 70% identity and 70% coverage for both sequences were clustered together using Markov Chain Clustering (MCL) (Inflation Value = 1.5) [130]. Proteins that did not pass these criteria with any other protein sequence in the HMM database were clustered separately into a new protein family. The protein sequences from the remaining 412 genomes were then iteratively scanned against the reference HMM database as described above, updating the HMM model and database after each iteration. Following the initial clustering of all proteins from the 413 *Pseudomonas* genomes into putative homolog families, HMM families were grouped into larger families if at least one member of a family shared more than 70% identity with at least one member of another family. Orthologous protein pairs were then extracted from these homolog families using the reciprocal pairwise distance approach and were clustered into ortholog families using MCL (Inflation Value = 1.5) [130].

Once all gene families had been clustered, we analyzed the pan-genome of *P. syringae* using a binary presence-absence matrix for each ortholog family in the 391 *P. syringae* genomes, where 1’s were used to encode presence and 0’s were used to encode absence [131]. We assigned all gene families that were present in at least 95% of the *P. syringae* strains in our dataset to the soft core genome and all other gene families to the accessory genome. The more lenient cutoff of 95% is justified because it allows us to limit the artificial reduction in the core genome size that occurs because of disrupted or unannotated core genes in some draft genomes (Figure S2). We then determined whether the pan-genome of *P. syringae* was opened or closed using the “micropan” R package [55]. Here, a rarefraction curve of the entire pan-genome was computed using 100 permutations, each of which was computed using a random genome input order. The curve was then fitted to Heap’s Law model to calculate the average number of unique ortholog clusters observed per genome and determine whether the pan-genome is opened or closed.

The phylogenetic relationships between the 391 *P. syringae* strains analyzed in this study were explored using both a soft core genome tree and a pan genome content tree. For the core genome tree, we multiple aligned the protein sequences from each soft core ortholog family using Kalign Version 2, which uses the Wu-Manber pattern matching algorithm [132]. We then concatenated these alignments and removed all monomorphic sites from this alignment using an in-house perl script. The core genome maximum likelihood phylogenetic tree was then constructed using FastTree with default parameters [133]. FastTree uses a combination of maximum likelihood nearest-neighbor interchange (NNIs) and minimum evolution subtree-pruning-regrafting (SPRs) methods for constructing phylogenies [133–135]. Local branch support values for the topology of the phylogenetic tree were also calculated in FastTree using Shimodaira-Hasegawa (SH) test [136]. For the genetic content tree, we used the shared gene-content information from the “micropan” R package to calculate the genetic distance between each strain and generate a pan-genome distance matrix with Jaccard’s method. The topological robustness of the gene content tree was tested by performing average linkage hierarchical clustering with 100 bootstraps. This same method was also employed for the effector content and EEL content trees.

### Identification and Analysis of Ecologically Relevant Genes

The first set of ecologically relevant genes that we investigated were the genes that constitute the T3SS, a key virulence determinant in pathogenic *P. syringae* strains. Specifically, we used the core structural genes of different forms of T3SSs, including the canonical tripartite pathogenicity island (T-PAI) T3SS, the atypical pathogenicity island (A-PAI) T3SS, the single pathogenicity island (S-PAI) T3SS, and the Rhizobium-like pathogenicity island (R-PAI) T3SS to explore the distribution of different T3SSs across the *P. syringae* species complex. To determine if a particular form of T3SS was present in a given strain, we performed a tblastn search for the core structural genes of each T3SS against each *P. syringae* genome assembly with an e-value cutoff of 1^−5^. All core structural genes for each T3SS were downloaded from NCBI GenBank, using *P. syringae* DC3000 and *P. viridiflava* PNA3.3a as references for the T-PAI T3SS, *P. syringae* Psy642 and *P. syringae* PsyUB246 as references for the A-PAI T3SS, *P. viridiflava* RMX3.1b as a reference for the S-PAI T3SS, and *P. syringae* 1448A as a reference for the R-PAI T3SS. We then chose the top hits for each T3SS structural gene in each genome, translated the region into a protein sequence, and confirmed that there were no premature truncations in the sequence. A given T3SS was considered present if all core structural genes for that T3SS were present and not truncated. These presence/absence data were then used to analyze the distribution of different T3SSs across the *P. syringae* species complex.

The second ecologically relevant genes that we explored were the T3SEs that are delivered into plant hosts by the T3SS. To analyze the distribution of T3SEs across the *P. syringae* species complex, we predicted known and novel T3SEs using discrete pipelines. For known T3SEs, we performed a tblastn against each *P. syringae* assembly using a collection of 1,215 experimentally verified or computationally predicted effector sequences downloaded from the BEAN 2.0 database (E-Value < 1^−5^) [75]. If a significant hit was identified for a T3SE, the region of the best or only hit was extracted from the genome as a putative T3SE. To identify novel T3SEs, we first constructed an HMM-model using known hrp-box motifs from three completely sequenced *P. syringae* genomes (*Pto* DC3000, *Pph* 1448A, and *Psy* B728A) [65, 70, 79, 137]. These motif sequences were multiple aligned using Kalign2 [132] and the HMM-model was constructed using hmmbuild [129]. The hrp-box HMM model was then scanned against each *P. syringae* genome assembly using nhmmer with a high e-value (10,000) and low bit score (4) threshold, given the likelihood that this model would yield false positives as a result of the short sequence length. Because a number of T3SEs are known to reside in operons, we then inspected the ten genes downstream of each predicted hrp-box motif for a N-terminal secretion signal using EffectiveT3 [138]. If a gene was both a less than 10 genes downstream of a hrp-box and classified as a T3SE based on their N-terminal secretion signal, we considered them putative novel T3SEs. The effector repertoire of each *P. syringae* strain was ultimately used to characterize the core and accessory effector profile of the *P. syringae* species complex.

A third set of ecologically relevant genes that we studied consisted of seven well-characterized phytotoxins of the *P. syringae* species complex, including coronatine, phaseolotoxin, tabtoxin, mangotoxin, syringolin, syringopectin, and auxin [76]. To determine if these pathways were present in each genome, we performed a tblastn search (E-Value < 1^−5^) using known proteins that are involved in the synthesis of each phytotoxin against each *P. syringae* genome assembly. Representative query sequences that are involved in the biosynthesis of each phytotoxin were obtained from GenBank, using strain PtoDC3000 for coronatine (17 biosynthesis genes), PsyBR2R for tabtoxin (20 biosynthesis genes), and PsyUMAF0158 for phaseolotoxin (17 biosynthesis genes), mangotoxin (10 biosynthesis genes), syringolin (6 biosynthesis genes), syringopectin (11 biosynthesis genes), and auxin (2 biosynthesis genes). If significant hits were found in a given genome for more than half the of the biosynthesis genes of a phytotoxin, it was considered present, and if not, the phytotoxin was considered absent. These presence/absence data were ultimately used to study the distribution of phytotoxins across the *P. syringae* species complex.

Finally, we also identified the complete collection of known virulence factors in each genome using the virulence factor database (VFDB, version R3), a reference database of bacterial protein sequences that contains more than 1,798 virulence factors from a total of 932 bacterial strains that represent 75 bacterial genera [63, 139, 140]. Specifically, we predicted virulence factors in each *P. syringae* genome by blasting the proteome of the genome against the entire VFDB (E-Value < 1^−5^). A protein sequence was considered a virulence factor if a hit was found that had more than 70% identity with a sequence in the VFDB database.

### Identification and Analysis of Evolutionarily Significant Genes

We classified any orthologous gene families that had one or more sites under positive selection as evolutionarily significant. To identify these ortholog families, we used the Fast Unconstrained Bayesian Approximation (FUBAR) pipeline to measure the ratio of non-synonymous substitution rates to synonymous substitution rates (*Ka/Ks*) at each site in each ortholog family [86]. The FUBAR pipeline was chosen because in implements a Markov Chain Monte Carlo (MCMC) sampler for inferring sites under positive selection, which makes it more efficient for inferring sites under positive selection in large alignments than other methods and allows us to account for the effects of recombination on signatures of selection [141]. For this analysis, we used the output of the GARD recombination analysis to partition ortholog families into non-recombinant fragments. We then analyzed both the partitioned and un-partitioned datasets using FUBAR with 10 MCMC chains, where the length of each chain was equal to 5,000,000, the burn-in was equal to 2,500,000, the Dirichlet Prior parameter was set to 0.1, and 1,000 samples were drawn from each chain. Evolutionarily significant genes were extracted from each genome if they were part of an orthologous family that had one or more sites under positive selection in the partitioned analysis.

### Detection of Genetic Recombination

We searched for signatures of homologous recombination within the *P. syringae* species complex using GARD [87], CONSEL [88], GENECONV [89], and PHIPACK [90] in all 17,807 ortholog families that were present in at least five strains. First, to generate input alignments for the recombination software, we independently aligned the nucleotide sequences for all ortholog families using translatorX [142], then heuristically removed sequences with a high frequency of gaps using the heuristic algorithm option (t=50) in MaxAlign [143]. For GARD, we analyzed the codon alignment of each family using default parameters, then parsed significant recombination breakpoints in the GARD results file. For CONSEL, we first constructed a protein tree and corresponding core genome tree for all strains in each ortholog family using FastTree [133]. CONSEL was then used with default settings to calculate and compare the per-site likelihood values for these two trees with the gamma option, and ortholog families that were significantly incongruent were identified as recombinant families. For GENECONV, we used a gscale parameter of 1 and otherwise default settings to detect significant signatures of recombination in each family based on the polymorphic sties in the multiple alignment. Lastly, for PHIPACK, we employed default settings to test for signatures of recombination based on the maximum chi-square (MaxChi2), the neighbor similarity score (NSS), and the pairwise homoplasy index (PHI) statistical frameworks [90]. The MaxChi2 method classifies ortholog families as recombining if a non-uniform distribution of sequence differences exists along the alignments. The NSS method classifies recombination when adjacent sites show significant incongruence compared to other sites. The PHI method computes an incompatibility score over a sliding window in the alignment using only parsimoniously informative sites, then calculates a p-value for recombination in the alignment by column permutation [90]. In all tests, recombination was considered significant if the p-value was less than 0.05 after correcting for multiple comparisons. Ortholog families with significant signatures of recombination in the GARD, CONSEL, GENECONV, and PHIPACK analyses were then combined to estimate recombination rates within the *P. syringae* species complex, after normalizing for the number of orthologs, the number of strains, and the branch lengths in each phylogroup. We also differentiated between intra- and inter-phylogroup recombination events for recombination events identified by GENECONV using their pairwise recombination rates.

In addition to assessing which gene families appear to be undergoing recombination within and between *P. syringae* phylogroups, we explored HGT between *P. syringae* and more distantly related species using a blastp search of all protein sequences in each *P. syringae* strain against the non-redundant NCBI GenBank database using an e-value cutoff of 1^−5^, a percent identity cutoff of 70%, and a percent query coverage cutoff of 70%. The top three blast hits were then extracted for each protein and the results were parsed to retain only matches from non-P. *syringae* species. Any of these remaining hits were viewed as potential HGT events. Although this approach is unlikely to provide accurate measures of the extent of HGT in the *P. syringae* species complex, it provides critical information on common donor and/or recipient species that may be sharing a niche and DNA with *P. syringae* strains.

### Estimating Relative Sequence Divergence (*Ka/Ks*)

For each *P. syringae* strain pair, we used the concatenated soft core genome alignments to calculate the pairwise rates of non-synonymous (*Ka*) and synonymous (*Ks*) substitution using the SeqinR package in R [59]. Average *Ka* and *Ks* values were then calculated for all phylogroups and between strains of different phylogroups. For comparison, we also calculated the evolutionary rates of a number of different distinct species pairs, including *A. hydrophila* (NC_0008570.1) – *A. salmonicida* (NC_009348.1, NC_004923.1, NC_004925.1, NC_004924.1, NC_009349.1, NC_009350.1), *N. gonorrhoeae* (NC_002946.2) – *N. meningitides* (NC_003112.2), *P. aeruginosa* (NC_002516.2) – *P. putida* (NC_009512.1), and *E. coli* (NC_002695.1, NC_002127.1, NC_002128.1) – S. *enterica* (NC_003198.1, NC_003384.1, NC_003385.1). Here, we identified core genes that were shared by each strain pair using a pairwise protein blast with an e-value threshold of 1^−5^, and sequence identity and query coverage cutoffs of 80%. We then aligned these core nucleotide sequences using TranslatorX and MUSCLE, and concatenated the alignments using a custom perl script. The *Ka* and *Ks* values for each of these species pairs were calculated using the SeqinR package in R, as we did with the *P. syringae* strains.

## TABLES

Not applicable.

## DATA ACCESS

All genomic data produced by this study have been submitted to NCBI. BioProject Accession numbers for all genomes sequenced in this study and all publically available genomes are available in Supplemental Dataset S1.

## ACKNOWLEDGMENTS

This work was supported by a Natural Sciences and Engineering Research Council of Canada Award to DSG and a Canada Research Chair in Comparative Genomics to DSG. We thank all members of the Guttman and Desveaux labs for helpful discussion and members of the CAGEF staff for technical support.

## AUTHOR CONTRIBUTIONS

S.T., D.G. designed the research; M.D., S.T., R.A., and D.G. analyzed the data; and M.D., S.T., and D.G. wrote the paper.

